# A Donor T-Cell Receptor Structural Signature Determines Alloreactive Potential and Predicts Acute Graft-Versus-Host Disease

**DOI:** 10.64898/2026.07.08.737087

**Authors:** Ximi K. Wang, Ajna Uzuni, Lingting Shi, David W. Harle, Rodney Macedo, Michael Pressler, Christian A. Gordillo, Kirubel Belay, Shami Chakrabarti, Joshua Fuller, Elizabeth O. Hexner, Alison W. Loren, David L. Porter, Markus Y. Mapara, Megan Sykes, Elham Azizi, Ran Reshef

**Affiliations:** Columbia Center for Translational Immunology, Department of Medicine, Columbia University Irving Medical Center, New York, New York, USA; Department of Biomedical Engineering, Columbia University, New York, NY, USA; Irving Institute for Cancer Dynamics, Columbia University, New York, NY, USA; Department of Computer Science, Columbia University, New York, NY, USA; Herbert Irving Comprehensive Cancer Center, Columbia University Irving Medical Center, New York; Blood and Marrow Transplantation and Cell Therapy Program, Division of Hematology & Oncology, Columbia University Irving Medical Center, New York, NY; Center for Cell Therapy and Transplant, University of Pennsylvania and Abramson Cancer Center, Philadelphia, PA

## Abstract

Acute graft-versus-host disease (GVHD) remains a lethal barrier to successful allogeneic hematopoietic cell transplantation, yet pre-transplant donor selection entirely ignores hypervariable T-cell receptor (TCR) architecture. Here, by characterizing over 64,000 alloreactive clonotypes, we demonstrate that human alloreactivity is dictated by a constrained, predictable baseline structural signature. Pathogenic, tissue-infiltrating alloreactive T cells exhibit significantly shortened CDR3β regions, altered antigen-facing biophysical features, biased VJ gene usage and extensive inter-donor sharing originating from public anti-pathogen memory reservoirs. These potent clones natively cluster within the high-frequency fraction of the unstimulated baseline donor repertoire. We introduce R50, an assay-independent metric quantifying this clonal dominance, which independently predicted a six-fold increased risk of acute GVHD in a cross-institutional cohort. This scalable *in silico* platform shifts pre-transplant risk stratification from HLA typing and demographic surrogates to precision immune-receptor modeling.

## Introduction

Allogeneic hematopoietic cell transplantation (HCT) is curative for various hematological malignancies and non-malignant conditions^1,2^. The development of acute graft-versus-host disease (GVHD) has remained a major barrier to a successful HCT, occurring in 23% to 52% of patients in contemporary cohorts and frequently progressing to life threatening or lethal complications^3,4^. As a result, there remains a need for identification of specific biomarkers, which would not only be able to robustly predict the development of GVHD, but also facilitate optimal donor selection prior to transplant. Importantly, most validated GVHD biomarker strategies are applied after transplantation and often after symptom onset, to risk-stratify outcomes and guide therapy, leaving a persistent gap in pre-transplant tools that can inform donor choice and prophylaxis strategy^5,6^.

Acute GVHD is mediated by donor derived T lymphocytes recognizing host antigens as foreign, activating adaptive immune pathways, resulting in activation of CD4 helper and CD8 cytotoxic T-cells, leading to destruction of host tissue primarily in the liver, skin, and gut^7,8^. These alloreactive T-cells constitute the key underlying mechanism of acute GVHD development^9,10^. Despite advances in GVHD prophylaxis and HLA matching algorithms, the incidence of acute GVHD remains high^11–13^, suggesting there is a need to uncover common features of alloreactivity within the pathogenic T lymphocyte population. A donor-derived measure of alloreactive potential could complement HLA matching and clinical risk factors, enabling more personalized donor selection. Identification of features unique to alloreactive T-cells would create diagnostic and therapeutic opportunities that could be exploited in GVHD and potentially in other settings such as solid organ transplants.

High-throughput T-cell receptor (TCR) sequencing of the third complementarity-determining region (CDR3) of the TCRβ chain, a key region for antigen specificity, has become a robust and established tool for dissecting the diversity and characteristics of T-cell populations^14–16^. Previous work has identified alloreactive T-cells in solid organ transplants by defining fingerprints of the alloreactive T-cell repertoire via *in vitro* mixed lymphocyte reactions (MLR)^17–19^. More recently, we adapted this strategy to the HCT setting and used it to identify alloreactive clones and track them post-transplant in the recipient over time^20^.

In our study, we enhanced the established MLR assay to uniquely profile alloreactivity in the HCT setting by incorporating recipient-derived professional antigen-presenting cells (APCs), which potentiates the detection of T-cell responses across all levels of HLA disparity, including in HLA-identical donor-recipient pairs. A patient-specific fingerprint of alloreactive T-cell sequences was identified via high throughput sequencing of TCRβ in the proliferating fraction of the MLR product^17,21^. We initially identified 64,146 unique alloreactive T-cell sequences in 20 donor-recipient pairs and characterized their features. Alloreactive T-cell clonotypes were similarly identified in tissue samples from the same patients at the time of suspected GVHD onset. We found that alloreactive T-cells are cross-reactive, frequently conserved between individual donors, and can be distinguished from nonalloreactive T-cells via differential CDR3β length and several biophysical parameters in the antigen-facing region. We then found that, while the overall cumulative frequency of alloreactive clones in the donor is low, they are typically expanded and aggregate in the top half of the donor repertoire in terms of individual frequencies. We then used the newly defined characteristics of alloreactive T-cell sequences to develop a predictive test for the development of GVHD based on the donor T-cell repertoire alone. Here, we identified R50, a T-cell population diversity metric with strong independent predictive value for acute GVHD based on donor T-cell composition pre-transplant, irrespective of HLA matching. Finally, we validated this metric in an expanded cohort of HCT donors from two institutions.

## Results

### Quantitative Landscapes of Donor Alloreactive T-cell Clonotypes

Using high-throughput TCRβ CDR3 sequencing, we identified alloreactive T-cell clonotypes using MLR assays enhanced with the addition of recipient-derived APCs that allows for greater sensitivity in 20 donor-recipient pairs^17,20,22,23^ (Discovery cohort; **Table 1**, **Fig. 1A)**. Clonotypes were classified as nonalloreactive if they were found exclusively in the donor pre-transplant or in the MLR-CFSElow (CFSElo) fraction but did not meet expansion criteria in the MLR (see Methods). A total of 64,146 unique alloreactive clonotypes and 1,567,045 nonalloreactive clonotypes were identified. For clinical validation of donor diversity metrics, we expanded the cohort by adding 37 transplant donors from 2 institutions, for whom CDR3β sequencing of pre-transplant donor samples was performed (Validation cohort; n = 57; **Supplemental Table 1).**

**Figure 1.**
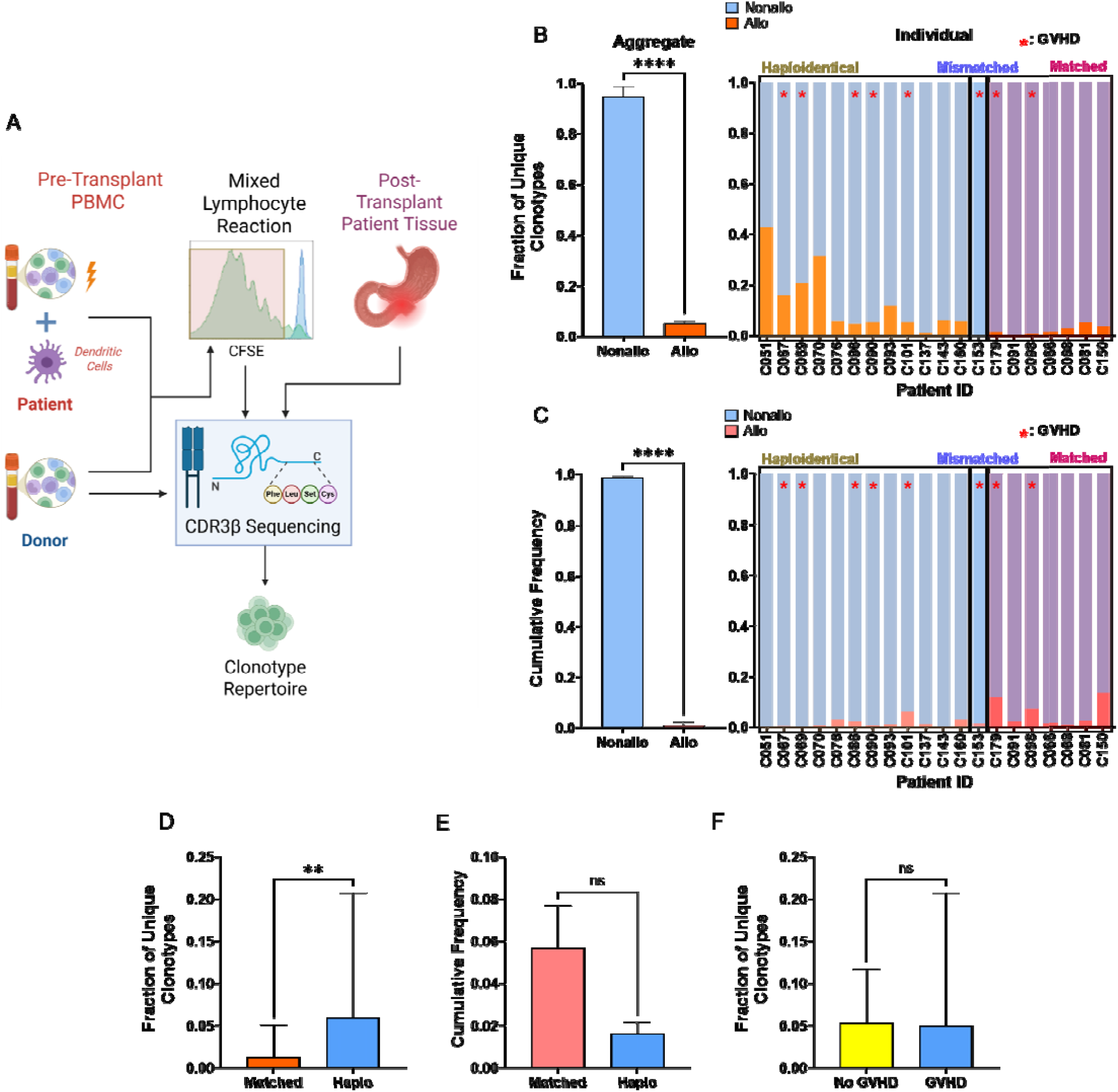
Mixed Lymphocyte Reaction (MLR) of Pre-transplant Donor and Recipient Peripheral Blood Mononuclear Cells (PBMC) Identifies Alloreactive Clonotypes as a Small Fraction of the Donor Repertoire. **A)** Experimental design: pre-transplant MLR and post-transplant tissue biopsies undergo bulk CDR3ꞵ sequencing. Both unstimulated donor T-cells and MLR-derived CFSE-low T-cells were sequenced. **B)** Comparison of the median fraction (95% CI) of unique alloreactive versus nonalloreactive clonotypes within the pre-transplant donor T-cell repertoire (Left: in aggregate; Right: individual donor-recipient pairs). Discovery cohort, n = 20, *p* < 1e^-15.^ Error bars depict the 95% confidence intervals. Red asterisk denotes patients who developed acute GVHD. **C)** Cumulative frequency of alloreactive vs nonalloreactive T-cells (Mean ±SEM) within the overall pre-transplant donor repertoire (Left: in aggregate, Right: in individual donors). Welch’s *t*-test used for comparison. **D)** Comparison of the median fraction (95% CI) of unique alloreactive clonotypes in HLA-mismatched haploidentical donors vs matched donors (combined matched sibling and unrelated), *p* = 0.0012. **E)** Cumulative alloreactive frequency (Mean ± SEM) of pre-transplant donor repertoire in matched (n = 12) vs haploidentical donor (n = 7), *p* = 0.0884. Welch’s *t*-test used for comparison. **F)** No association between acute GVHD (grade 2-4) and the fraction of unique alloreactive clonotypes (median with 95% CI) in the donor T-cell repertoire, *p* = 0.4727. Mann-Whitney U test used for median comparisons.

**Table 1.**
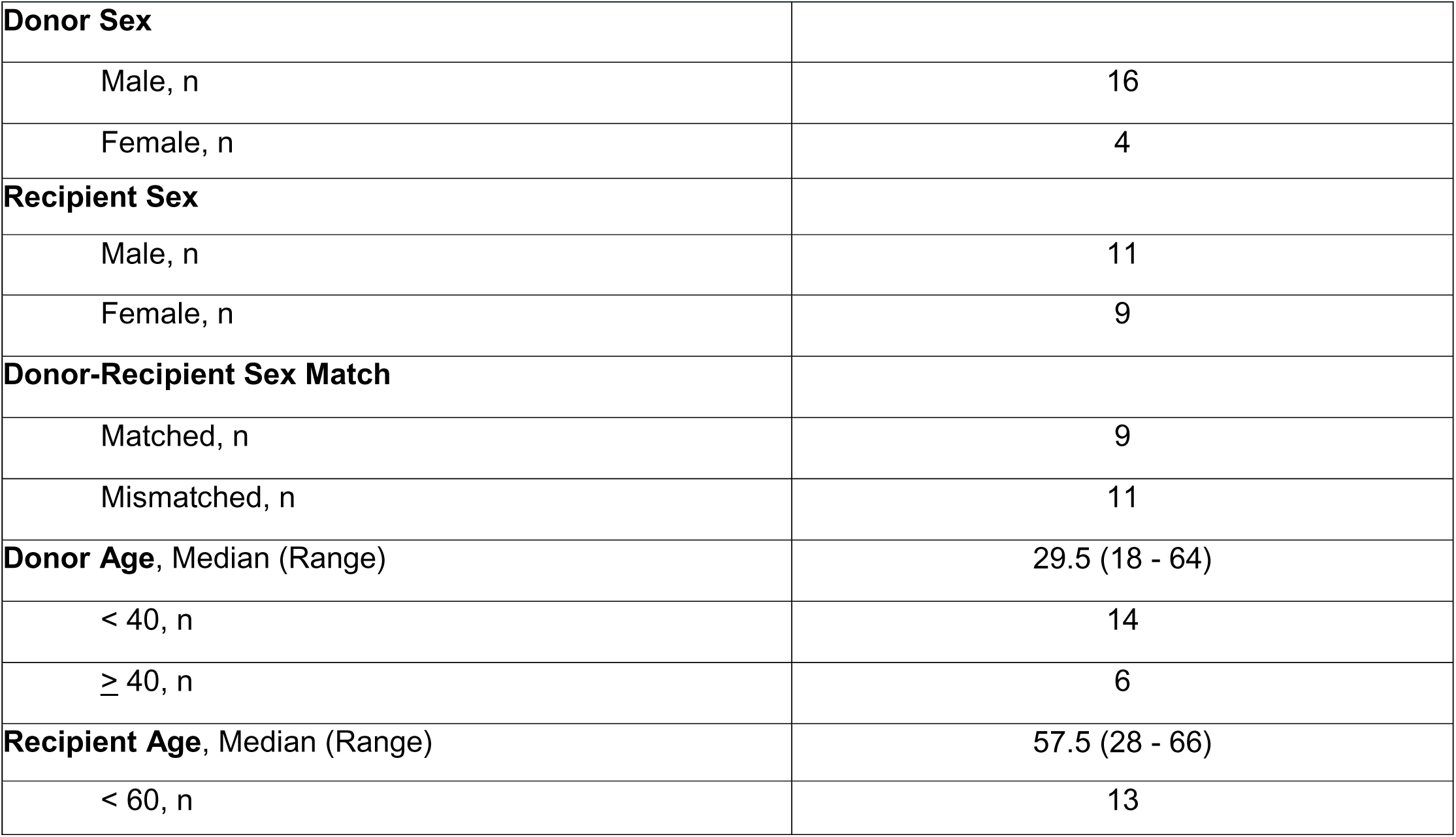
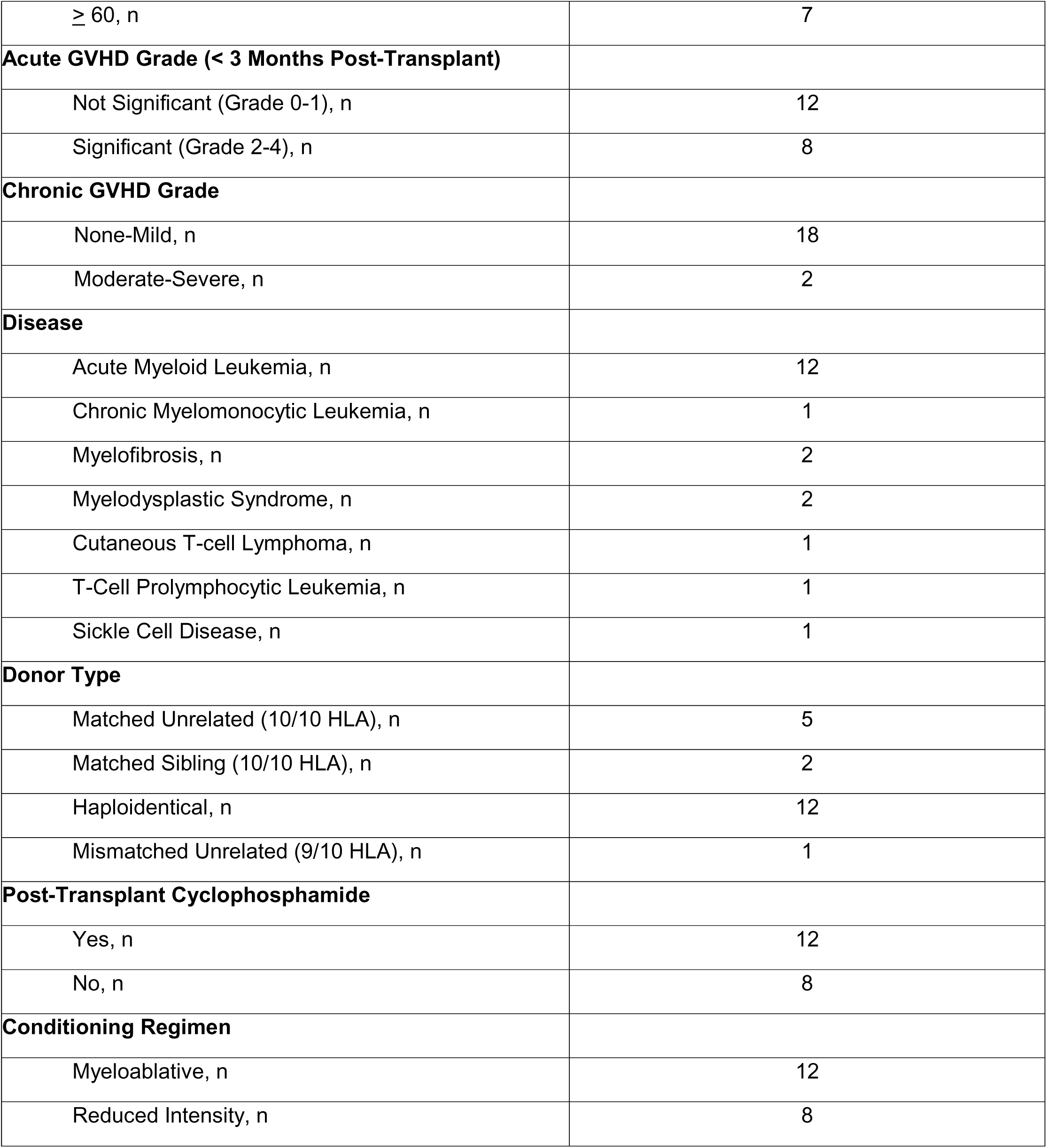
Patient and Donor Characteristics in the Discovery Cohort (n = 20)

Consistent with previously described results^21^, we found that alloreactive clonotypes are a small fraction of the baseline donor repertoire, whether measured by unique clonotype count or total T-cell frequency. In terms of unique TCR diversity, alloreactive clonotypes represented a small fraction of all unique donor-derived T lymphocyte clonotypes compared to nonalloreactive (*p* < 1e^-10^) with a median of 0.0521 (range 0.0027 to 0.4264) among the 20 discovery cohort pairs **(Fig. 1B)**. This was mirrored at the quantitative level within the total T-cell pool; the mean cumulative frequency of all alloreactive T-cells within pre-transplant donor repertoire was 0.0279 (range 0.0004 to 0.1336, vs. nonalloreactive *p* < 1e^-15^) **(Fig. 1C)**. The fact that alloreactivity is driven by genetic disparity was reflected in the fraction of unique alloreactive clonotypes, which was significantly higher in haploidentical donors compared to HLA-matched donors (median 0.0596 vs 0.0130, *p* = 0.0012). **(Fig. 1D);** it was not reflected in either the absolute number of unique alloreactive clonotypes **(Supplemental Fig. 1A)** or their cumulative frequency within the entire baseline donor repertoire (mean alloreactive frequency 0.0571 vs 0.0164, *p* = ns) **(Fig. 1E)**. Interestingly, neither the number of alloreactive clonotypes **(Supplemental Fig. 1B)**, the mean cumulative frequency of all alloreactive T-cells **(Supplemental Fig. 1C)**, nor the fraction of unique alloreactive clonotypes in the donor correlated with clinically significant acute GVHD post-transplant **(Fig. 1F)**. There was also no difference in the fraction of unique alloreactive clonotypes by other stratifications including matched unrelated vs matched sibling donors, CMV+ vs CMV- donors, younger vs older donors, male vs female donors, or donors with recipient sex mismatch vs sex matched **(Supplemental Fig. 1D)**.

### Alloreactive T-cells Exhibit Shorter CDR3β Regions

We previously defined expansion patterns of individual alloreactive clones over time post-transplant in HCT recipients using the *DecompTCR* computational tool. In this analysis, we identified that clonotypes that followed patterns associated with severe GVHD were characterized by shorter CDR3β and greater hydrophobicity compared to patterns associated with mild or no GVHD^20^. Here, we wanted to examine whether similar features more generally distinguish alloreactive from nonalloreactive clones. To this end, we conducted a pooled analysis of unique alloreactive clonotypes in comparison to nonalloreactive clonotypes across the 20-discovery cohort donor-recipient pairs. A comparison of the CDR3β length distributions revealed that alloreactive clonotypes were significantly shorter than their nonalloreactive counterparts **(Fig. 2A)**, and that alloreactive CD4^+^ clonotypes were also shorter in comparison to CD8^+^ clonotypes **(Fig. 2B).** We did not have information regarding CD4^+^ and CD8^+^ T-cells separately within the nonalloreactive T-cell population.

**Figure 2.**
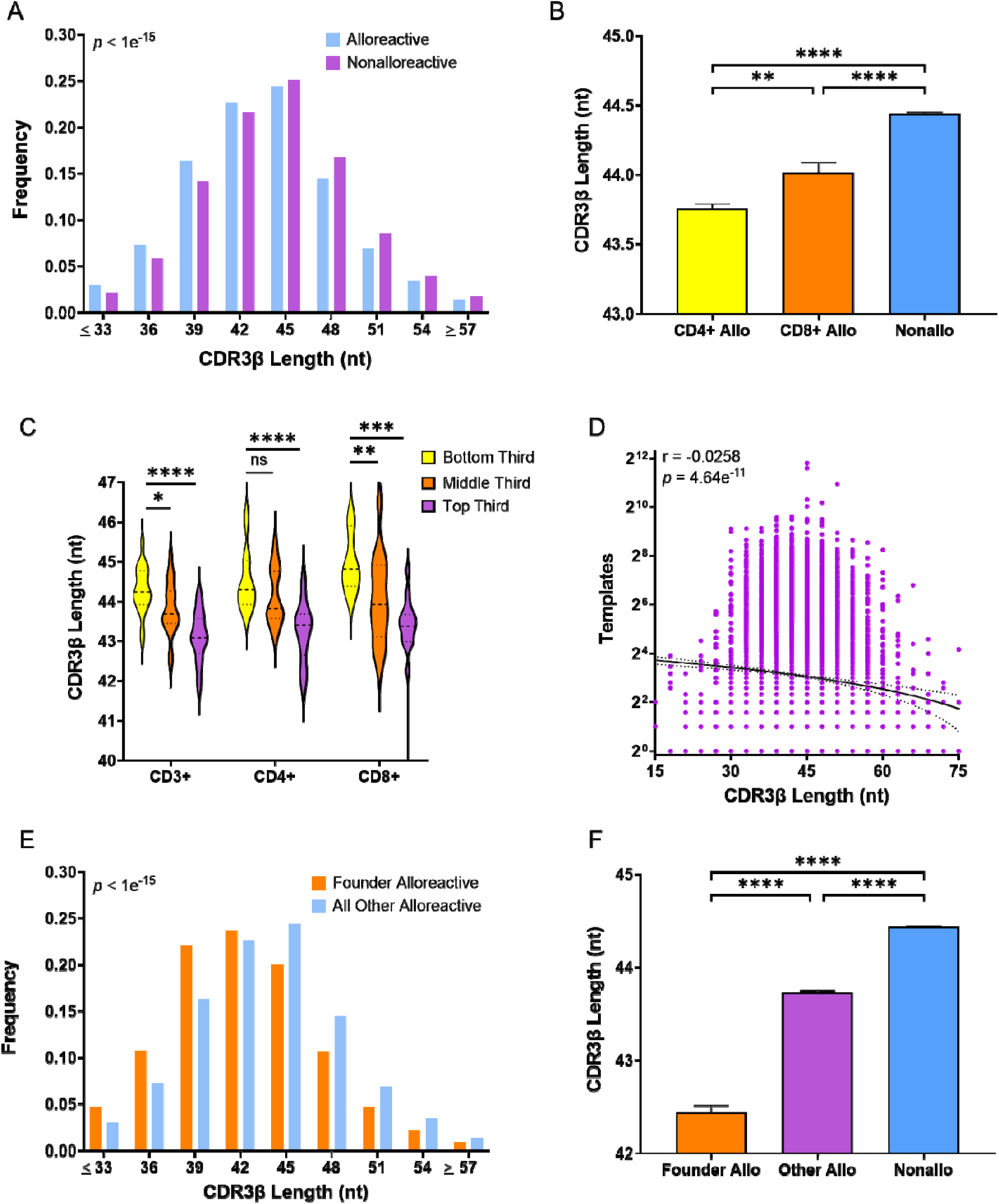
Alloreactive Clonotypes Have Shorter CDR3ꞵ Length in Comparison to Nonalloreactive Clonotypes. **A)** Distribution of CDR3β length (number of nucleotides; nt) among pooled unique alloreactive versus nonalloreactive clonotypes (p < 1e^-15^; Welch’s *t*-test). **B)** Comparison of mean CDR3β length in CD4^+^ and CD8^+^ alloreactive vs all nonalloreactive clonotypes. **C)** Violin plots comparing the distributions of CDR3β length of alloreactive clonotypes stratified into tertiles based on their clonal abundance (template counts) within the CFSElo fraction, comparing total CD3^+^, CD4^+^, and CD8^+^ alloreactive clonotypes. **D)** Shorter CDR3ꞵ length in alloreactive clonotypes correlates with significantly increased abundance of these clones in the proliferating fraction of the MLR (CFSElo). Pearson correlation coefficient calculated to establish significance. **E)** Comparison of CDR3β length among all alloreactive vs founder alloreactive clonotypes (defined as measurable in the donor). **F)** Mean CDR3β length distribution of founder alloreactive, non-founder alloreactive, and nonalloreactive clonotypes. \**p* <0.05, \*\**p* < 0.01, \*\*\**p* < 0.001, \*\*\*\**p* < 0.0001. Welch’s *t*-test used for all mean comparisons. Error bars reflect SEM.

We investigated whether specific donor characteristics could account for differences in overall donor CDR3β length. Our analysis revealed that mean CDR3β length of unique donor repertoire clonotypes was not significantly influenced by HLA matching, donor sex, or age group. Furthermore, there was no direct correlation between donor age and CDR3β length **(Supplemental Fig. 2A-D)**. These findings suggest that shorter CDR3β length is a defining characteristic of alloreactive clonotypes, independent of other donor characteristics.

### Shorter CDR3β Correlates with in vitro Clonal Expansion and in vivo Repertoire Abundance of Alloreactive Clones

We then sought to identify specific functional attributes of alloreactive clonotypes that are associated with this shortening of CDR3β length. First, we evaluated clonal proliferation within the MLR by stratifying the overall alloreactive T-cell population by their frequency within the CFSElo repertoire. The top third group, which represents the most frequent clonotypes, had significantly shorter CDR3β length compared to the bottom third. This held true across both CD4^+^ and CD8^+^ subsets **(Fig 2C)**. Similarly, when classifying alloreactive clonotypes into expanded (> 2 templates) and unexpanded (≤ 2 templates) in the CFSElo fraction, the expanded clonotypes exhibited significantly shorter CDR3β regions **(Supplemental Fig. 3A)**. The significant negative correlation between CDR3β length and clonal expansion in the MLR suggests that shorter CDR3β sequences are structurally linked to a higher degree of alloreactivity **(Fig. 2D)**.

Given that part of our objective was to develop a clinically relevant metric that could be applied to the baseline donor repertoire without requiring a MLR, we next examined whether the structural hallmark of alloreactive clones also characterized the alloreactive clones that were detectable in the donor. Because many MLR-identified alloreactive clones exist at frequencies too low to be detected in baseline donor blood samples, we defined “Founder” alloreactive clonotypes as those present in sufficient quantities to be measurable within the unstimulated donor repertoire. Crucially, these founder clonotypes exhibited significantly shorter CDR3β length when compared to either nonalloreactive **(Supplemental Fig. 3B)** or even non-founder alloreactive clonotypes **(Fig. 2E-F)**. When stratified into frequency tertiles in the CFSElo, founder alloreactive clonotypes maintained a significantly shorter CDR3β in comparison to non-founder alloreactive clonotypes **(Supplemental Fig. 3C).** In conclusion, shorter CDR3β sequences distinguish the most expanded alloreactive clonotypes across both the *in vitro* CFSElo proliferation fraction and the baseline donor repertoire. The fact that these structurally constrained clones dominated both compartments implies that reduced CDR3β length marked a population with a profound, intrinsic alloreactive potency. Consequently, profiling this distinct structural feature provides a scalable strategy to map the repertoire of high-risk donor clonotypes directly from an unstimulated pre-transplant blood draw.

### Tissue-Infiltrating Alloreactive (TIA) Clonotypes Are Clinically Relevant, Measurable in The Donor and Exhibit a Short CDR3β Signature

To determine whether MLR-defined alloreactive clonotypes are clinically relevant mediators of disease, we tracked their presence within GVHD target tissues (gastrointestinal tract and skin) post-transplant. We sequenced the TCRβ repertoire from post-transplant tissue biopsies and matching peripheral blood mononuclear cells (PBMCs) from 7 HCT recipients at the onset of suspected GVHD symptoms (**Supplemental Table 2)**. This analysis successfully identified 183 MLR-defined alloreactive clonotypes actively infiltrating affected tissues.

These tissue-infiltrating alloreactive (TIA) clonotypes were detected in corresponding PBMC samples at a significantly higher frequency than tissue-infiltrating nonalloreactive (TIN) clonotypes (27.8% vs. 14.9%), indicating their involvement in a systemic process **(Fig. 3A, Supplemental Fig. 4A)**. Crucially, these clinically relevant TIA clonotypes exhibited the defining molecular signature of alloreactivity: their CDR3β sequences were significantly shorter than both TIN clonotypes and the broader MLR-defined alloreactive population **(Fig. 3B-C)**.

**Figure 3.**
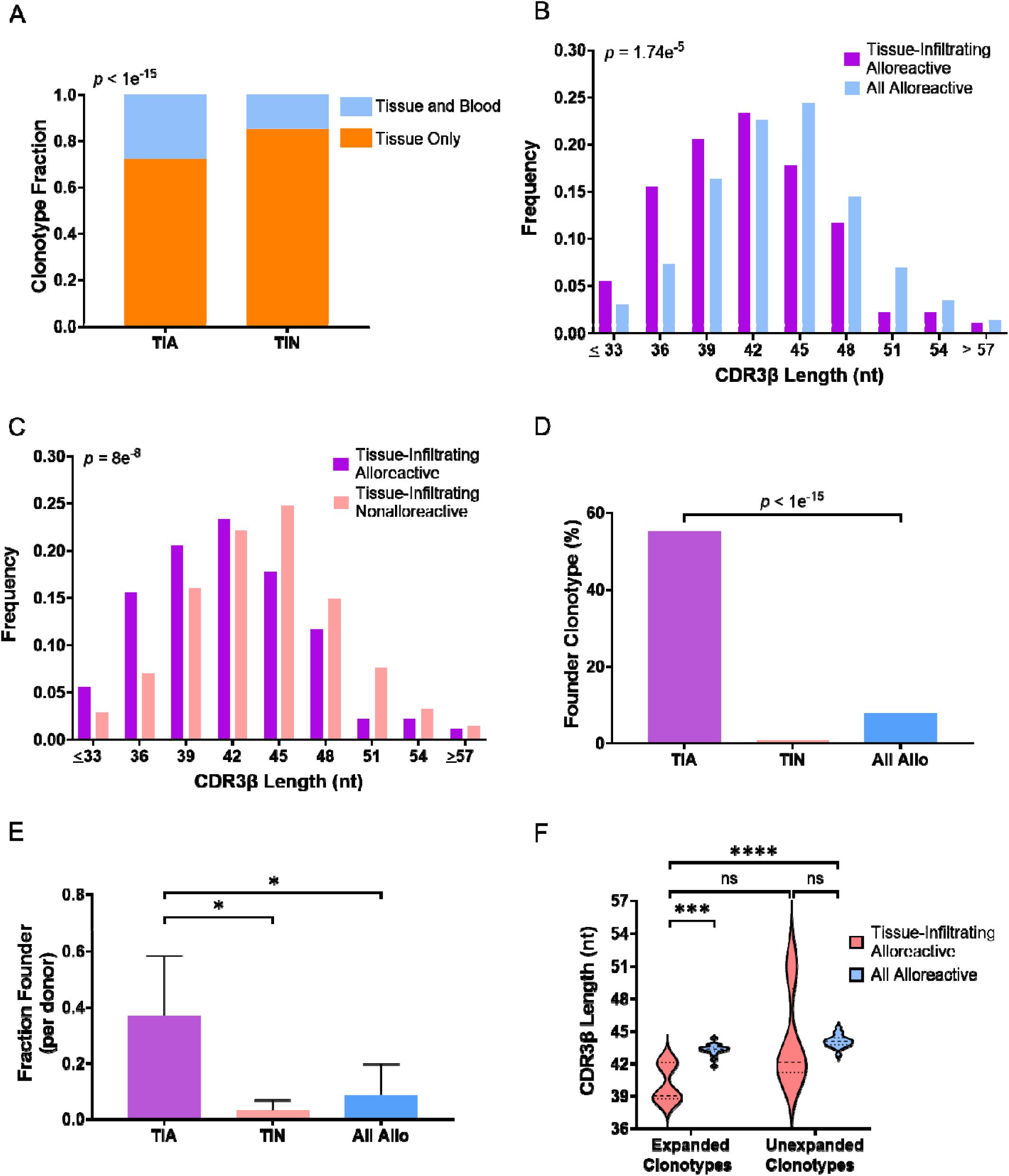
Tissue-infiltrating Alloreactive Clonotypes Have the Overall Shortest CDR3β and Are Commonly Founder Clones. **A)** Bar plots showing the percentage of unique tissue-infiltrating alloreactive (TIA) and tissue-infiltrating nonalloreactive (TIN) clonotypes that are restricted exclusively to recipient target tissues (gut or skin) versus those concurrently detectable in peripheral blood. TIA clones are significantly more likely to circulate systemically (identifiable in both tissue and blood) than TIN clones (27.8% vs. 14.9%; OR 2.21, 95% CI 1.61 – 3.03, p < 1e^-15^). **B)** Comparison of CDR3β length distributions between tissue-infiltrating alloreactive (TIA) clonotypes versus all pooled alloreactive clonotypes in 7 patients with matching gut or skin biopsies at the time of suspected GVHD onset. **C)** Comparison of CDR3β length distributions between TIA clonotypes versus TIN clonotypes. **D)** Percentage of total unique clonotypes detectable in pre-transplant donor repertoire, shown across all donors. TIA clonotypes are more likely to be founder alloreactive than all alloreactive clonotypes (OR 14.63, 95% CI 10.97 - 19.70, *p* < 1e^-15^) **E)** Median fraction of clonotypes from each category for HCT with post-transplant tissue biopsies that were founder clonotypes (detectable in the pre-transplant donor repertoire) per donor. Mann-Whitney U test used for comparisons, error bars depict the 95% confidence intervals, n = 7. **F)** Violin plots comparing CDR3β length within expanded (> 2 templates in MLR-derived CFSElo fraction) and unexpanded (≤ 2 templates) subpopulations of TIA clonotypes against all pooled alloreactive clonotypes. \**p* < 0.05, \*\**p* <0.01, \*\*\**p* < 0.001, \*\*\*\**p* < 0.0001. Welch’s *t*-test used for all comparisons of means.

Furthermore, the TIA clonotypes involved in GVHD *in vivo* were predominantly founder clones, reinforcing the potential of the pre-transplant donor repertoire as a biomarker. More than half (55.2%) of TIA clonotypes were readily detectable in the unstimulated pre-transplant donor sample, compared to only 7.77% of the overall alloreactive population **(Fig. 3D)**. The proportion of founder clones was significantly higher among TIA clonotypes than among TIN clonotypes (median 0.40 vs. 0.011, *p* = 0.039) or all alloreactive clonotypes (median 0.40 vs. 0.0521, *p* = 0.027) **(Fig. 3E)**. Finally, TIA clones that expanded significantly in the MLR-CFSElo fraction possessed shorter CDR3β sequences than all other alloreactive clones **(Fig. 3F).** This observation further solidifies the link between reduced CDR3β length, robust *in vitro* expansion, and *in vivo* pathogenic potential.

### Alloreactive T-cells are Drawn from a Public Repertoire of Shared and Cross-reactive Clonotypes

The presence of “public” T-cell clonotypes shared between individuals with a high degree of cross reactivity has been previously demonstrated in mechanistic studies done in humanized mice and linked to shorter CDR3β length^24^. Whether this pool of shared, cross-reactive T-cells constitutes the primary source of clinical alloreactivity in human HCT has remained an open question. Given our finding that alloreactive clonotypes have shorter CDR3β regions, we hypothesized that they would be significantly enriched for publicly shared sequences. We evaluated this by defining a clonotype as “shared” if it appeared in more than one donor in our discovery cohort. Of the total 1,631,191 unique clonotypes in our dataset, 114,026 were shared (6.99%). In contrast, of the 64,146 MLR-defined unique alloreactive clonotypes, 12,539 (19.55%) were shared, demonstrating that alloreactive clonotypes were > 3 times more likely to be public than nonalloreactive clonotypes (OR 3.10, 95% CI 3.04 - 3.16) **(Fig. 4A)**. A majority of the shared clonotypes in the dataset (64.2%) are found in only 2 distinct donors **(Supplemental Fig. 4B)**.

**Figure 4.**
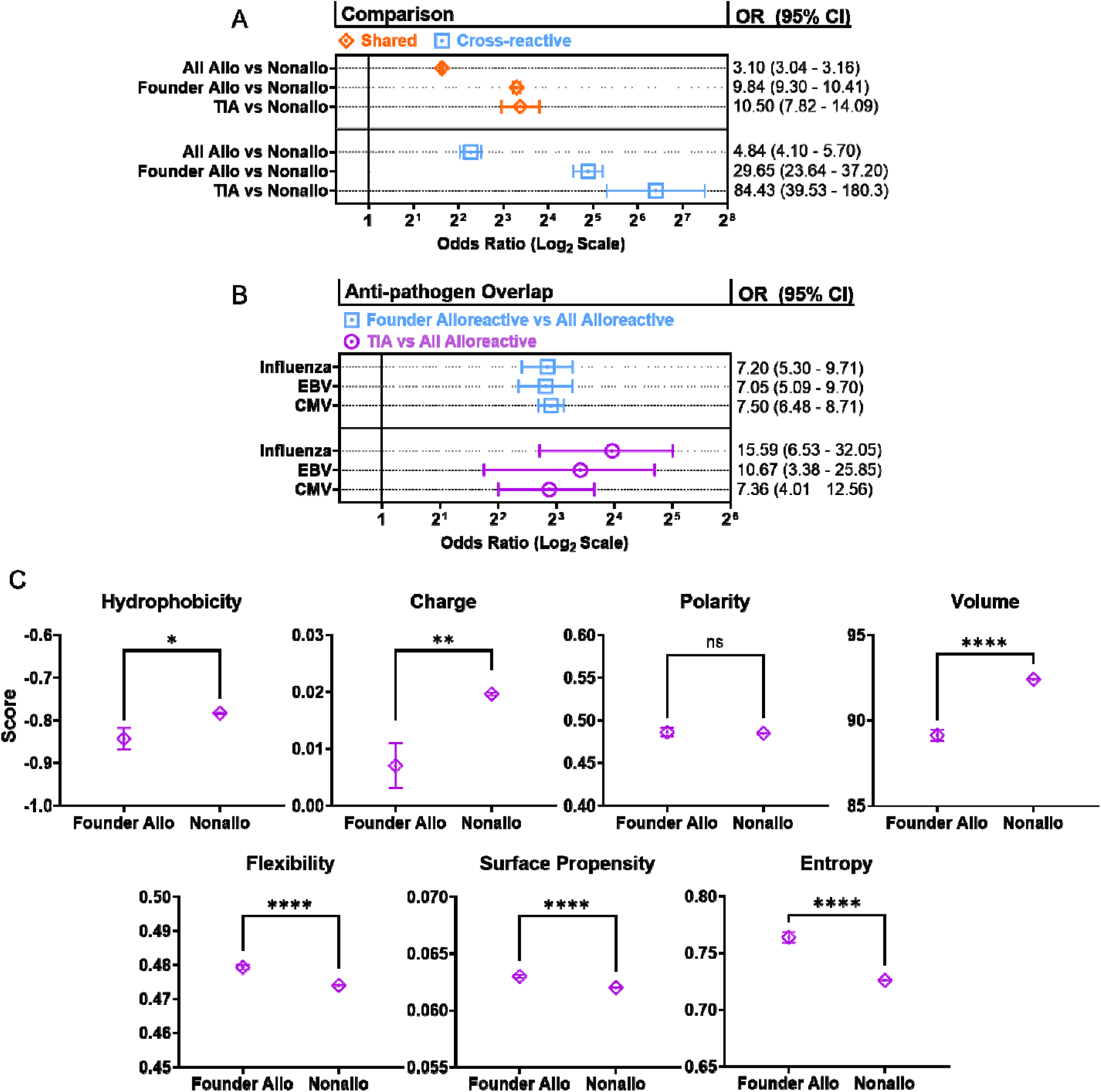
Highly Alloreactive Clonotypes are Shared and Cross-reactive, Overlap with Anti-pathogen Clonotypes, and Have Distinct Biochemical Features at Key CDR3β Positions. **A)** Odds ratios (95% CI) of the likelihood of alloreactive clonotypes to be shared or cross-reactive compared to nonalloreactive clonotypes. Tissue-infiltrating (TIA), founder alloreactive, and all alloreactive clones analyzed separately. **B)** Odds ratios (95% CI) for enrichment of known anti-pathogen sequences (CMV, EBV, Influenza A/B) among TIA and founder alloreactive clonotypes relative to all alloreactive clonotypes. **C)** Calculated means (±SEM) of hydrophobicity (Kyte-Doolittle scale), charge, polarity, volume, flexibility, surface propensity, and entropy for amino acid residues in position 6 and position 7 of CDR3β chain for founder alloreactive vs nonalloreactive clonotype groups. Statistical significance is based on FDR-adjusted *p*-values (Benjamini–Hochberg), with *q* < 0.05 considered significant; \**q* < 0.05, \*\**q* < 0.01, \*\*\**q* < 0.001, \*\*\*\**q* < 0.0001. Welch’s *t*-test used for all comparisons unless otherwise specified.

This enrichment was even more pronounced in the founder clonotypes and the tissue-infiltrating clonotypes that are more functionally relevant to clinical disease. 43.54% of founder alloreactive clonotypes and 42.08% of the TIA clonotypes from GVHD target tissues were shared **(Supplemental Fig. 4C)** and exhibited substantially higher odds of being shared compared to nonalloreactive clonotypes (Founder: OR 9.84, 95% CI 9.30 - 10.41. TIA: OR 10.50, 95% CI 7.82 - 14.09) **(Fig. 4A).**

Building on the established association between sharedness and cross-reactivity, we then examined whether our MLR-defined alloreactive T-cells are cross-reactive. We compared the alloreactive clonotypes to a dataset of 3,797 previously identified cross-reactive CDR3β sequences, defined as clonotypes interacting with multiple unique epitopes^24,25^. We found that alloreactive clonotypes had significantly higher odds of being cross-reactive than their nonalloreactive counterparts (OR 4.84, 95% CI 4.10 - 5.70). Furthermore, this effect was more pronounced in founder clonotypes (OR 29.65, 95% CI 23.64 - 37.20) and most in TIA clonotypes (OR 84.43, 95% CI 39.53 - 180.3), which suggests a very high likelihood of cross-reactivity among the pathogenic alloreactive clones **(Fig. 4A).**

### Alloreactive Clonotypes Overlap with a Public Repertoire of Anti-Pathogen T-Cells

A prevailing hypothesis in transplantation immunology suggests that memory T-cells generated during prior pathogen exposures can cross-react with alloantigens, forming a pre-existing reservoir of alloreactive cells^26^. Our findings that alloreactive clonotypes have shorter CDR3β and are highly shared align closely with the known characteristics of these "public" anti-pathogen T-cell populations. To directly test this link, we analyzed publicly available TCRβ repertoires specific for common pathogens, including Epstein-Barr Virus (EBV), Cytomegalovirus (CMV), and Influenza A/B.

First, we confirmed that these anti-pathogen clonotypes share the key molecular signature of alloreactivity; their CDR3β length was similar to alloreactive clonotypes and was significantly shorter than nonalloreactive clonotypes **(Supplemental Fig. 5A)**. These anti-pathogen clonotypes are also exceptionally public, being 8 times more likely to be shared - CMV (OR 8.48, 95% CI 8.12 - 8.86), EBV (OR 8.42, 95% CI 7.65 - 9.24), and Influenza (OR 8.33, 95% CI 7.65 - 9.06) in comparison to nonalloreactive and > 2 times more likely to be shared than alloreactive clonotypes **(Supplemental Fig. 5B)**.

Importantly, we found clear functional overlap between anti-pathogen and alloreactive responses. Clonotypes specific to CMV, EBV and Influenza were all significantly more likely to be found in the MLR-defined alloreactive pool than in the nonalloreactive pool (OR > 3 for all viruses; **Supplemental Fig. 5C)**. This link to clinical GVHD was strongest in both founder alloreactive clonotypes and in the TIA clonotypes identified in GVHD biopsies. Founder alloreactive clonotypes (OR > 7 for all viruses) and TIA clonotypes were far more likely to match known anti-pathogen clonotypes than the broader alloreactive repertoire: CMV (OR 7.36, 95% CI 4.01 - 12.56), EBV (OR 10.67, 95% CI 3.38 - 25.85), and Influenza (OR 15.59, 95% CI 6.53 - 32.05), supporting the conclusion that highly alloreactive clonotypes are associated with increased cross-reactivity and may represent memory T cells that originate from prior exposure to pathogen-derived alloantigens **(Fig. 4B).**

### Highly Alloreactive Clonotypes Exhibit Specific Biochemical Features at Key Antigen-Contacting Sites

In humanized mice, shared TCR sequences have been shown to exhibit reduced hydrophobicity and enrichment of neutral or hydrophilic amino acids at key antigen-facing positions (positions 6 and 7) of the CDR3β chain^24^. Distinct biochemical features, most notably hydrophobicity, strongly influence reactivity and thymic T-cell selection^27–29^. Because we established that founder clones represent the most potent, highly expanded, and clinically relevant (tissue-infiltrating) subset of alloreactive T-cells detectable in the pre-transplant donor, we specifically evaluated their average charge, polarity, volume, surface propensity, conformational entropy, flexibility, and hydrophobicity at these critical positions using established metrics^30–32^. We found that, compared to nonalloreactive clonotypes, the CDR3β sequences of founder alloreactive clonotypes are significantly less hydrophobic, more negatively charged, smaller in volume, more flexible, display greater surface propensity, and have greater entropy at positions 6 and 7 (**Fig. 4C)**. Notably, these biochemical distinctions are less pronounced when analyzing the broader, unstratified alloreactive pool, reinforcing that these specific biophysical signatures strongly correlate with the highest degree of alloreactivity (i.e., the highest capacity for *in vitro* expansion and tissue infiltration) (**Supplemental Fig. 6A**). Consistent with these findings, founder alloreactive clonotypes display a bias toward altered amino acid composition at CDR3β positions 6 and 7 relative to the broader alloreactive and nonalloreactive repertoire **(Supplemental Fig. 6B)**, statistically confirmed by per-position chi-squared analysis of amino acid composition.

### Biased VJ Gene Usage Serves as a Biomarker for Alloreactivity and GVHD

Building on the finding that shorter CDR3β sequences define alloreactive T cells, we examined underlying V(D)J recombination features and found that MLR-defined alloreactive clonotypes demonstrated reduced nucleotide insertions at the V–D and D–J junctions **(Supplemental Fig. 6C)**. We next investigated whether the flanking V and J gene segments that frame the CDR3β region exhibit usage biases correlating with MLR-defined alloreactivity and clinical GVHD risk^33,34^. We compared VJ pairing frequencies across MLR-defined alloreactive versus nonalloreactive clonotypes **(Fig. 5A)**, as well as between unstimulated donor repertoires from donors whose recipients did (GVHD+) or did not (GVHD-) develop grade 2-4 acute GVHD within 3 months post-transplant **(Fig. 5B).** This analysis revealed three VJ pairings significantly enriched in both alloreactive and GVHD+ clonotype populations (termed “GVHD+/Allo-biased”; *n* = 23,855 unique clonotypes). Conversely, eight pairings demonstrated the opposite bias, being significantly overrepresented in nonalloreactive clones and GVHD- repertoires (termed “GVHD-/Nonallo-biased”; *n* = 41,228 unique clonotypes).

**Figure 5.**
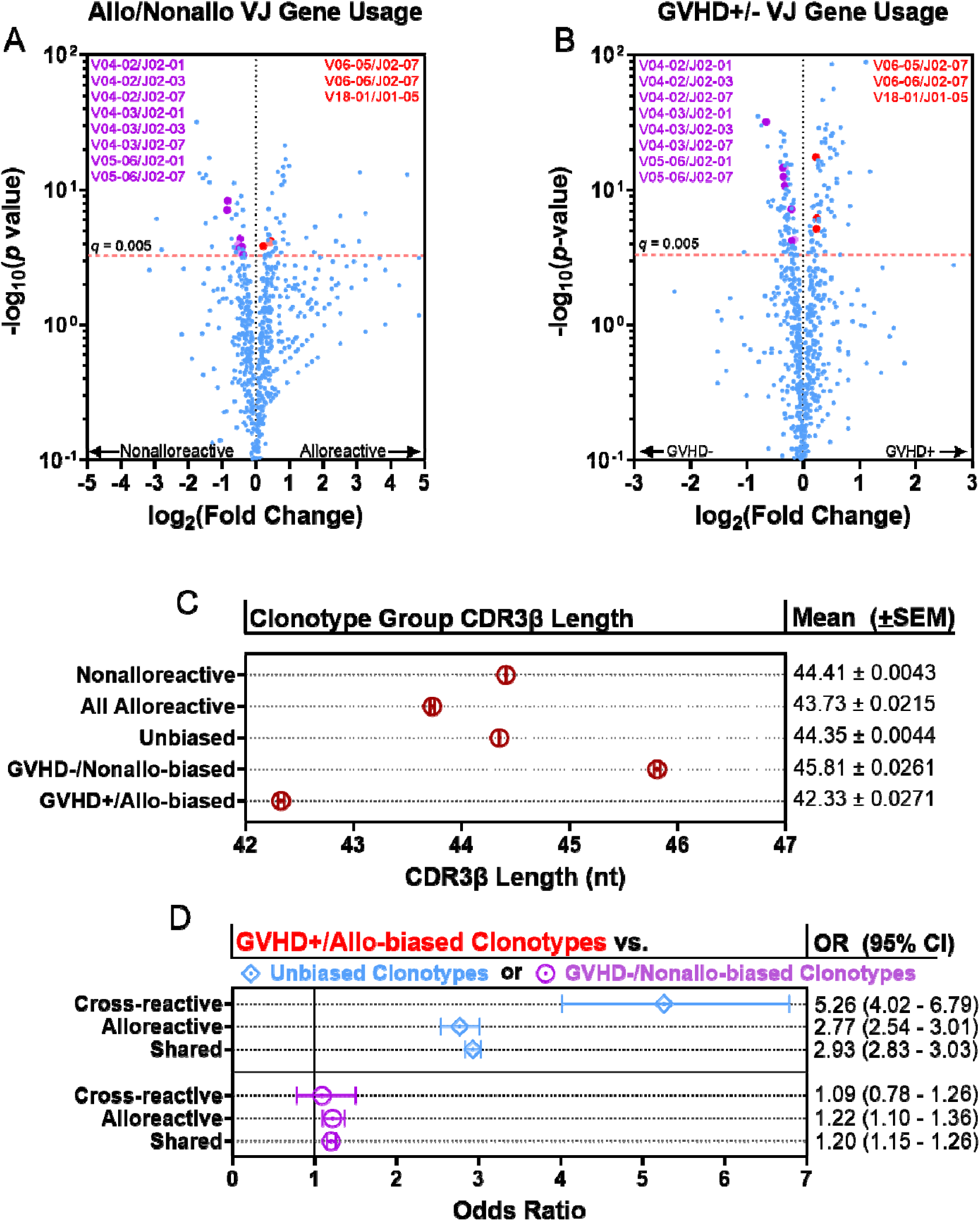
VJ Gene Usage of Alloreactive and GVHD-Associated Clonotypes. Volcano plots showing VJ gene combination usage in **A)** alloreactive vs. nonalloreactive clonotypes and **B)** clonotypes from donors whose recipients did (GVHD+) or did not (GVHD-) develop grade 2-4 acute GVHD. Fisher’s exact test used to calculate the significance of the VJ frequency between the groups. Benjamini-Hochberg test with a *q* value of 0.005 as the cut-off used to determine statistical significance. Alloreactive VJ pairings that significantly correlated with GVHD are highlighted. Red values denote VJ pairings that are both significantly GVHD+ and Alloreactive biased (GVHD+/Allo-biased VJ pairings) and purple values denote a significant association with GVHD- and Nonalloreactive (GVHD-/Nonallo-biased VJ pairings). Blue represents Unbiased VJ pairings. **C)** Comparison of mean CDR3β lengths (±SEM) of unique GVHD+/Allo-biased clonotypes to GVHD-/Nonallo-biased clonotypes, clonotypes from all other VJ pairings (Unbiased), all MLR-derived alloreactive clonotypes, and all nonalloreactive clonotypes. Welch’s *t*-test used for all. *p* < 0.0001 for all comparisons. **D)** Odds ratio (95% CI) of the GVHD+/Allo-biased clonotypes being cross-reactive, shared, and alloreactive in comparison to GVHD-/Nonallo-biased and all Other Clonotypes.

We next profiled the CDR3β length and repertoire features of these distinct subpopulations of clonotypes. Consistent with our earlier findings, GVHD+/Allo-biased clonotypes exhibited significantly shorter CDR3β regions than both their GVHD-/Nonallo-biased (mean 42.33 vs 45.81, *p* < 1e^-15^) and Unbiased counterparts (mean 42.33 vs 44.35, *p* < 1e^-15^). Furthermore, GVHD-/Nonallo-biased clonotypes had significantly longer CDR3β than Unbiased Clonotypes (45.81 vs 44.35, *p* < 1e^-15^), aligning with a reduced alloreactive preference. Finally, the GVHD+/Allo-biased population was highly enriched for alloreactive, shared and cross-reactive clonotypes. Notably, GVHD+/Allo-biased clonotypes demonstrated a significantly higher probability of inter-donor sharing (OR 1.20, 95% CI 1.15 - 1.26) and alloreactivity (OR 1.22, 95% CI 1.15 - 1.26) than the GVHD-/Nonallo-biased group **(Fig. 5D).**

### Alloreactive Clonotypes Cluster in the High-Frequency Fraction of the Donor Repertoire and Correlate with GVHD

Given that shared clonotypes often exhibit high frequencies within overall T-cell repertoires^35^, we hypothesized that the alloreactive and shared clonotypes we identified would be disproportionately expanded in the baseline donor repertoire, and that this abundance would correlate with acute GVHD. To map their distribution, we segmented each baseline donor’s repertoire into deciles based on clonal frequencies, ranging from the most abundant clones contained in the lowest-diversity deciles to the least abundant clones in the highest diversity deciles **(Fig 6A)**.

**Figure 6.**
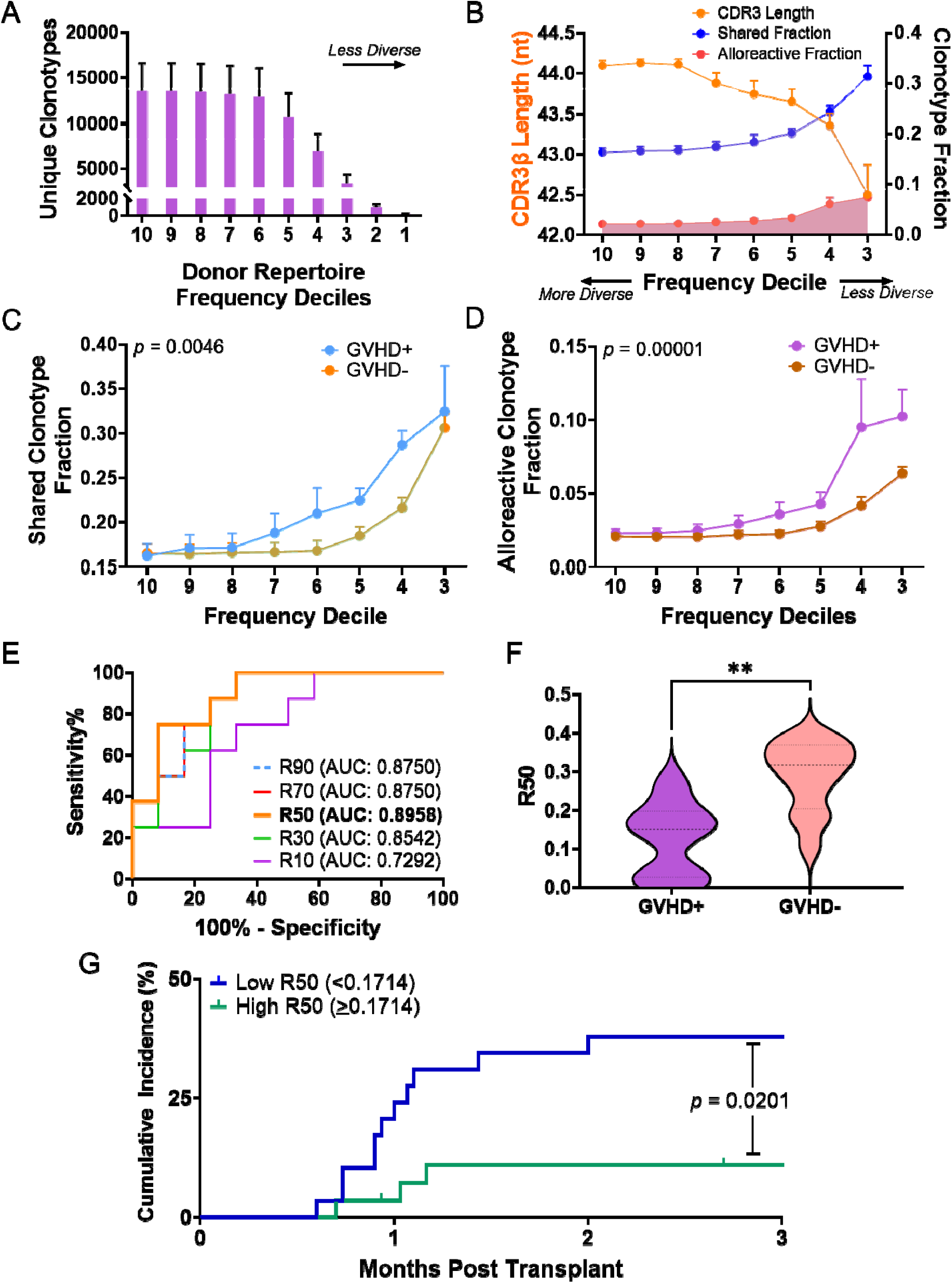
Alloreactive Clonotypes Cluster in the High-Abundance Fraction of the Donor Repertoire, and Low R50 Diversity Independently Predicts Acute GVHD. **A)** Mean number (±SEM) of unique clonotypes per frequency decile with decreasing diversity. **B)** Mean unique clonotype CDR3β length, fraction of shared clonotypes, and fraction of alloreactive clonotypes within the donor T-cell repertoire are plotted for each decile of clonal frequencies. First and second deciles are not presented given that in some donors the top clonotypes had frequencies >10%. **C)** Comparison of mean (± SEM) shared clonotype fraction in deciles for patients with and without acute GVHD in the discovery cohort. 2-way ANOVA test used to determine statistical significance. **D)** Comparison of mean (±SEM) alloreactive clonotype fraction by decile frequency intervals for patients with and without GVHD. 2-way ANOVA test used to determine statistical significance. **E)** Receiver Operating Characteristics (ROC) curve for prediction of acute GVHD using the diversity of the donor repertoire above decile XX (RXX). **F)** Median donor R50 (± 95% CI) for patients with and without acute GVHD. Mann-Whitney U test used to determine significance. **G)** Cumulative incidence plots showing time to acute GVHD for the expanded validation cohort (n=57), separated using the derived optimal diagnostic cutoff for R50. Mantel-Cox test used for comparison.

We found that shared and alloreactive clonotypes are not randomly distributed in the donor repertoire; rather, they are significantly enriched within the high-frequency deciles. This enrichment coincided with a progressive decrease in average CDR3β length, revealing a clear inflection point near the 5^th^ decile (**Fig. 6B)**. Crucially, the composition of this high-frequency fraction of the donor repertoire correlated with clinical outcomes. In donors whose recipients subsequently developed acute GVHD, this fraction exhibited significantly shorter CDR3β length **(Supplemental Fig. 7A)** and a greater representation of both shared **(Fig. 6C)** and alloreactive **(Fig. 6D)** clonotypes. These findings strongly suggest that the clonal architecture of the top half of a donor’s T-cell repertoire is a key determinant of GVHD risk and a viable target for a predictive biomarker.

### Low Donor R50 Diversity is an Independent Predictor of Acute GVHD

Directly identifying alloreactive T-cells requires complex *in vitro* functional assays (MLRs) that are impractical for routine pre-transplant screening. However, having established that potent alloreactive clonotypes natively concentrate and dominate the high-frequency fraction (top half) of the unstimulated donor repertoire, we reasoned that the overall diversity of this specific fraction could serve as a readily measurable, assay-independent surrogate for alloreactive potential. A repertoire heavily skewed by these expanded, public clones would present as having lower diversity specifically within its top frequency deciles.

To test this, we adapted a previously described TCR diversity measurement^21,22^, evaluating RXX - a metric that quantifies T-cell diversity strictly within the top XX% of a repertoire’s most abundant clones. Using a Receiver Operating Characteristic (ROC) analysis in the discovery cohort (n = 20), we evaluated various RXX thresholds for predicting acute GVHD. Aligning with the biological inflection point identified earlier, R50 - representing the diversity of the top 50% of clones - emerged as the most predictive metric (AUC 89.58%, PPV 79.42%, NPV 89.54%, *p* = 0.0034), outperforming other RXX thresholds (**Fig. 6E; Supplemental Table 3)**. We established an optimal R50 cutoff of 0.1714 to distinguish between high- and low-diversity donors. Applying this cutoff, donors linked to acute GVHD development exhibited significantly lower R50 diversity compared to the no-GVHD group (median 0.1504 vs 0.3160, *p* = 0.0022) **(Fig. 6F)**. In contrast, acute GVHD was not predicted by assessing overall donor repertoire diversity using conventional metrics, including the Shannon index, Simpson’s clonality, and Simpson’s index **(Supplemental Fig. 7B).**

Applying this cutoff, low R50 donors in the discovery cohort demonstrated a strikingly higher 3-month cumulative incidence of acute GVHD compared to high R50 donors (88.89% vs. 14.88%, *p* = 0.0003) **(Supplemental Fig. 7C)**. To validate this finding, we evaluated the same cutoff in an expanded, cross-institutional cohort (n = 57; **Supplemental Table 2)**. The prognostic association proved robust: low R50 donors were again linked to a significantly elevated cumulative incidence of acute GVHD (37.93% vs. 10.99%, *p* = 0.0201) **(Fig. 6G)** at 3-month post-transplant, and median R50 values remained distinct between the GVHD and no-GVHD groups (0.1357 vs. 0.1962, *p* = 0.0434) **(Supplemental Fig. 7D).**

To verify that R50 reflects the underlying biology of alloreactivity, we analyzed its correlates within the discovery cohort. Repertoires from donors with low R50 scores were significantly skewed toward the high-risk, GVHD+/Allo-biased VJ pairings (2 of 3), whereas high R50 repertoires favored the protective, GVHD-/Nonallo-biased pairings (6 of 8) **(Supplemental Fig. 8A).** Furthermore, the high-frequency fraction of low R50 repertoires was characterized by significantly shorter CDR3β sequences and a greater proportion of shared clonotypes **(Supplemental Fig. 8B-D).** These features biologically anchor the mathematically-derived R50 metric to the structural hallmarks of pathogenic alloreactivity. This predictive value appears distinctly tied to the initial alloreactive process, as R50 was not significantly associated with chronic GVHD at any time post-transplant **(Supplemental Fig. 8E)**.

Finally, we evaluated the independent clinical utility of tested R50 against established risk factors in the expanded cohort. In univariate analysis, a low R50 score significantly predicted acute GVHD (HR 4.059, *p* = 0.0170) **(Supplemental Table 4)**. Critically, in a multivariable Cox regression model adjusting for donor and recipient age, sex mismatch, post-transplant cyclophosphamide (PTCy) use, HLA mismatch, disease type, and donor CMV status, R50 emerged as the sole significant independent predictor. Patients receiving grafts from low R50 donors faced a > 6-fold increased risk of developing acute GVHD (HR 6.37, 95% CI: 1.57 - 33.60; *p* = 0.0087) **(Supplemental Table 5)**. We found no association between R50 and disease relapse or overall survival **(Supplemental Tables 6 and 7)**, underscoring its specificity as a pre-transplant biomarker for acute GVHD risk.

## Discussion

We demonstrate that human alloreactive T-cells possess reproducible structural and repertoire-level features that distinguish them from nonalloreactive T-cells. Consistent with previously described results, alloreactive T-cells constitute a small numerically minor fraction of the entire donor repertoire^21^. Interestingly, while major HLA-mismatch significantly expands the repertoire breadth (the fraction of unique alloreactive clonotypes), it does not quantitatively alter their overall cellular mass (cumulative frequency) in the donor. This decoupling suggests that while initial genetic disparity determines the structural diversity of the alloreactive pool, post-transplant allorecognition and GVHD prophylaxis regimens dictate the actual survival and selective elimination of these clones^36^. Notably, by leveraging recipient antigen-presenting cells (APC)-enhanced MLRs in clinically relevant HCT pairs, we provide the first direct evidence that human alloreactive T-cells are fundamentally characterized by significantly shorter CDR3β sequences than their nonalloreactive counterparts.

The foundational premise of our findings addresses a critical gap in current pre-transplant risk assessment: while contemporary donor selection algorithms rely almost exclusively on HLA matching and broad demographic surrogates^37^, they fundamentally fail to account for the hyper-variable, highly individualized architectural variations native to donor TCR repertoires. By demonstrating that human alloreactivity is dictated by constrained, predictable baseline structural sequence signatures and clonal frequency distributions, our study offers a paradigm shift toward predictive, repertoire-based risk stratification. Rather than depending on cumbersome, low-throughput *ex vivo* cellular assays (i.e., MLRs), we show that clinical alloreactive potential can be computationally forecasted using an accessible metric like R50. This metric successfully captures macro-repertoire clonal distributions directly from baseline donor sequencing data prior to graft collection. Crucially, the clinical utility of this quantitative framework extends far beyond HCT. By establishing a scalable *in silico* platform to decode immune receptor architectures, these insights provide generalizable modeling paradigms that can be adapted to predict allograft rejection risk in solid organ transplantation and optimize the precision design of next-generation allogeneic cellular therapies.

The existence of publicly shared^38,39^, cross-reactive T-cells demonstrate that unique selective pressures shape the human alloreactive repertoire. These shared structural features of alloreactive T-cells likely extend beyond HCT; cross-reactive memory T-cells shaped by identical thymic and peripheral selection pressures are implicated in donor-reactive responses in solid organ transplantation^40^, indicating that CDR3β-based repertoire profiling may have broad utility in predicting allograft rejection. At the point of origin within the thymus, these shared clonotypes arise from shorter, less hydrophobic CDR3β sequences that facilitate passage through positive selection while evading negative selection^24^. However, interpreting the absolute abundance of these clones must account for peripheral selection, which serves as the primary driver of their expansion in human blood^41^. Because short CDR3β lengths inherently confer high cross-reactivity, these clones maintain an increased likelihood of expanding in response to microbial antigens encountered throughout life.

This history of antigen exposure heavily shapes the pool of donor T-cells poised to mediate acute GVHD^42^. As these cross-reactive clones expand against pathogens, they transition into the memory T-cell compartment, vastly increasing their peripheral frequency relative to naïve T-cells^43^. This peripheral expansion explains the prominence of "founder" alloreactive clones: they are readily detectable in unstimulated, pre-transplant populations precisely because they are highly abundant memory cells. As previously indicated by DeWolf and Sykes^21^, while both naïve and memory compartments contribute to the alloresponse, these abundant memory cells are far more likely to be captured in baseline sampling and exhibit the dominant expansion in MLRs required to be defined as alloreactive. While peripheral blood reflects only a limited portion of the tissue-resident T-cell repertoire after HCT^44^, these abundant circulating memory clones are precisely the population captured at baseline, and consequently, the clinical acute GVHD response in target tissues is heavily seeded by this pre-existing, cross-reactive memory reservoir.

Importantly, these human-centric observations directly challenge historical murine data asserting that effector and memory T-cells lack the capacity of inducing acute GVHD^45,46^. Although these preclinical models inspired clinical translation through clinical trials utilizing naïve T-cell depletion to prevent GVHD^47,48^, those trials ultimately focused primarily on preventing *chronic* GVHD, rather than acute disease. Furthermore, randomized controlled studies to validate this concept are lacking. By longitudinally tracing alloreactive clones from baseline donor blood into inflamed post-transplant tissue biopsies, our findings provide human validation that overrides the preclinical dogma, re-establishing the critical pathogenic role of donor memory T-cells as mediators of human acute GVHD.

The discovery that the alloresponse is significantly shaped by a public, pre-existing pool of cross-reactive memory T-cells forms the biological basis for our R50 metric. Because these high-risk clones predictably cluster in the high-frequency fraction of the donor repertoire, quantifying the diversity of the top 50% most abundant clones (R50) captures a donor’s alloreactive potential. Broadened to a cross-institutional cohort, R50 served as a robust, independent predictor of acute GVHD, even when adjusting for known clinical variables such as donor age and PTCy use^4,49,50^. While PTCy improves GVHD-free survival by decreasing post-transplant repertoire diversity and removing potent allogeneic clonotypes^51–53^, it frequently fails to eliminate all pathogenic clones^20^, leaving the baseline risk captured by R50 highly relevant.

Our study is limited by the clinical constraints of acquiring pre-transplant donor-recipient PBMC for high-volume MLRs, as well as our exclusive sequencing of the TCRβ chain, though CDR3α is known to play a lesser role in antigen recognition^54^ and identification of clonality using TCRβ alone is accurate in >95% of cases^20^. Additionally, while R50 strongly predicts acute GVHD, it did not predict chronic GVHD in our cohort, warranting further studies with increased statistical power for late-onset complications.

In conclusion, human alloreactive T lymphocytes are distinguishable by distinct CDR3β lengths, biochemical features, and VJ gene pairing biases. Shaped by both thymic constraint and peripheral, microbe-driven memory expansion, these cross-reactive clones serve as the principal mediators of the acute GVHD response. Translating these fundamental immunologic insights into the R50 diversity metric provides a powerful, assay-independent method to predict acute GVHD. Integrating baseline TCR repertoire profiling into clinical algorithms has the potential to profoundly improve donor selection and enable risk-adapted prophylaxis in allogeneic HCT.

## Materials and Methods

### Patient Inclusion Criteria

Donor/Recipient pairs for the discovery cohort were patients with hematological diseases undergoing HCT at Columbia University Medical Center in New York, NY, USA. This cohort was in part previously presented and published^20^. A representative mix of donor types, conditioning regimens and GVHD prophylaxis regimens was selected with MLR experimentation conducted **(Table 1)**. For validation of the R50 metric, a cohort of 34 transplant donors and recipients treated at the University of Pennsylvania and 3 additional Columbia University donors and recipients were added **(Supplemental Tables 1)**. Sample sizes reflect the number of HCT recipients and donor-recipient pairs available within the IRB-approved clinical protocols at Columbia University and the University of Pennsylvania.

### Acute Graft-versus Host Disease (GVHD) definition

GVHD in this study is classified as acute GVHD grade 2-4, *i.e.* clinically significant GVHD in accordance with MAGIC guidelines^55^, within 3-months post-transplant.

### Sample Acquisition

Recipient and donor blood samples were acquired from HCT donors and recipients at baseline prior to transplant. Post-transplant recipient blood samples and tissue biopsies from lower GI tract, upper GI tract and skin were collected at the time of suspected clinical diagnosis of GVHD. Blood samples were processed via a Ficoll density gradient to obtain PBMCs and were cryopreserved per standard laboratory protocol.

### Monocyte-derived Dendritic Cell Maturation

The addition of recipient-derived dendritic cells increases the sensitivity of the MLR and yields more cell divisions in the CFSElo portion^23^. Matured dendritic cells were thus derived from CD14^+^ cells^56^.

Cryopreserved recipient PBMCs were thawed, washed, and resuspended in FACS buffer. CD14 MicroBeads (CD14 MicroBeads, human, Miltenyi Biotec, catalog no. 130-050-201) were used to positively select CD14^+^ cells, according to the manufacturer’s protocol.

CD14^+^ cells were resuspended in sterile R10 media (RPMI supplemented with 10% human serum, 0.6% HEPES, 10mg/mL Gentamycin, and 200mmol/L Glutamine) at 1 x 10^6^ cells/mL per well in a 24-well flat bottom plate. Granulocyte-macrophage colony-stimulating factor (R&D Systems Human GM-CSF Recombinant Protein, Fisher Scientific, catalog no. 215GM010CF) and IL-4 (R&D Systems Human IL-4 Recombinant Protein, Fisher Scientific, catalog no. 204IL010CF) were added to each well at concentrations of 50ng/mL and 20ng/mL, respectively. The CD14^+^ cell cultures were incubated at 37°C for 9 days. On day 7, wells were supplemented with TNF-⍺ (TNF-⍺ human, Millipore Sigma, catalog no. SRP3177-50UG) and Prostaglandin E2 (Prostaglandin E2, Fisher Scientific, catalog no. 22-961-0) at concentrations of 25ng/mL and 1 ug/mL, respectively. Cells were harvested at the end of day 9, washed, and resuspended in R10 media at 2 x 10^6^ cells/mL.

### Mixed Lymphocyte Reaction

The mixed lymphocyte reaction protocol was adapted from previously described methods^17^. Cryopreserved responder (donor) PBMCs were thawed, washed, resuspended in 1xPBS at 1 x 10^6^ cells/mL, and stained with CFSE at a concentration of 0.05 micromolar (CellTrace CFSE Proliferation Kit, Thermo Fisher Scientific, catalog no. C34554). CFSE-stained cells were incubated for 20 minutes at 37°C, washed once with R10 media, and resuspended in R10 media at 2 x 10^6^ cells/mL.

Cryopreserved stimulator (recipient) PBMCs were thawed, washed, resuspended in 1xPBS at 1 x 10^6^ cells/mL, and stained with Far Red at a concentration of 0.01 micromolar (CellTrace Far Red Cell Proliferation Kit, Thermo Fisher Scientific, catalog no. C34564). Far Red-stained cells were incubated for 20 minutes at 37°C, washed once with R10 media, resuspended in R10 media at 6 x 10^6^ cells/mL, and irradiated at 35 Gy.

100uL (200,000 cells, in total) of CFSE-stained responders, 100uL (600,000 cells, in total) of Far Red-stained stimulators, and 20uL (40,000 cells, in total) of monocyte derived dendritic cells were plated in each well of a 96 well round bottom plate, keeping the concentration of responders: stimulators: dendritic cells at 5:15:1. Subsequently, MLR cultures were incubated at 37°C for six days.

### MLR and Donor Flow Cytometry

On day 6 of culture, MLR cells were harvested, washed, and resuspended in FACS buffer. Cell surface staining was performed for 30 minutes at 4°C with fluorochrome-conjugated antibodies against CD19 (BD clone HIB19, catalog no. 561295), CD14 (BD clone MΦP-9, catalog no. 562693), CD56 (BD clone B159, catalog no. 560842), CD3 (BD clone UCHT-1, catalog no. 555335), CD4 (BD clone RPA-T4, catalog no. 560769), and CD8 (BD clone SK1, catalog no. 560273). Cells were then washed, resuspended in FACS buffer, and stained with cell viability solution (Via-Probe Cell Viability Solution, BD, catalog no. 555815). FACS sorting was performed on a BD Influx cell sorter. In order to obtain the donor-reactive populations, depending on the quantity of cells present, cells were either sorted for CD19^-^CD14^-^CD56^-^Far-Red^-^CFSE^lo^CD3^+^ or CD19^-^CD14^-^CD56^-^Far-Red^-^CFSE^lo^CD3^+^CD4^+^ and CD19^-^CD14^-^CD56^-^Far-Red^-^CFSE^lo^CD3^+^CD8^+^.

Cryopreserved unstimulated pre-transplant donor and post-transplant recipient PBMCs were also thawed, washed, and resuspended in FACS buffer. Cell surface staining was performed for 30 minutes at 4°C with fluorochrome-conjugated antibodies against CD19 (BD clone HIB19, catalog no. 561295), CD14 (BD clone MΦP-9, catalog no. 562693), CD3 (BD clone UCHT-1, catalog no. 555335), CD4 (BD clone RPA-T4, catalog no. 560769), and CD8 (BD clone SK1, catalog no. 560273). Cells were then washed, resuspended in FACS buffer, and stained with cell viability solution (Via-Probe Cell Viability Solution, BD, catalog no. 555815). Depending on the quantity of cells present, cells were either sorted for CD19^-^CD14^-^CD3^+^ or CD19^-^CD14^-^CD3^+^CD4^+^ and CD19^-^CD14^-^CD3^+^CD8^+^.

### DNA Isolation and TCRβ Sequencing

Genomic DNA was isolated from post-transplant biopsy specimens using the Qiagen DNeasy Blood & Tissue Kit (catalog no. 69504), per manufacturer’s protocol and stored at -20°C.

Genomic DNA was isolated from cell populations in the MLR, donor, and recipient post-transplant PBMC samples using the Qiagen QIAamp DNA Mini Kit (catalog no. 51304), per manufacturer’s protocol and stored at -20°C. DNA was shipped on dry ice to Adaptive Biotechnologies for TCRB sequencing. Datasets were downloaded from the Adaptive ImmunoSEQ Analyzer 3.0 at the amino acid level, restricted to productive rearrangements, and reported as productive frequency with no minimum frequency threshold applied.

### Identifying Circulating and Alloreactive T-cell Repertoires

Datasets were downloaded as described above and the templates identified in the CFSElo MLR samples were compared to the corresponding unstimulated donor samples. Unique T-cell clonotypes were identified based on their CDR3β amino acid (*aa*) sequences, which indicate the biochemical properties of antigen recognition^24^. Individual alloreactive T-cells were identified from CFSElo repertoires using previously described criteria^17,22^. A clonotype was classified as alloreactive if its CFSElo frequency exceeded 1.99-fold that of the unstimulated donor (rather than a strict 2-fold cutoff, to capture borderline clonotypes in low-frequency repertoires), or if it was absent from the donor repertoire but present in the CFSElo fraction at a frequency of > 2e^-5^. CD4^+^ and CD8^+^ CFSElo clonotypes were compared with CD4^+^ and CD8^+^ donor samples when possible. If not available, they were compared with CD3^+^ donor samples. If only CD3^+^ CFSElo samples were available, they were compared with CD3^+^ donors only. In total, 64,146 unique alloreactive clonotypes and 1,567,045 nonalloreactive clonotypes were produced from all MLRs donor-recipient pairings because of this analysis. Alloreactive clonotypes identified in post-transplant tissue biopsies were defined as tissue-infiltrating alloreactive clonotypes (TIA). Nonalloreactive clonotypes identified in post-transplant tissue biopsies were defined as tissue-infiltrating nonalloreactive clonotypes (TIN). Founder alloreactive clonotypes were defined as alloreactive clonotypes with detectable signal in the pre-transplant unstimulated donor repertoire.

### Measurement of Repertoire Diversity

R50 score was chosen as the diversity measurement of pre-transplant unstimulated donor clonotype repertoire, which measures diversity for the 50% of the TCR CDR3β population with the most abundant individual clonotypes, indicating the extent to which this population is expanded in contrast to the bottom 50%. This is obtained by sorting the unique *aa* clonotypes in decreasing order starting with the highest frequency sequence and going descending order. *N(T_50_)* is the least number of unique clonotypes which constitute 50% of total repertoire.

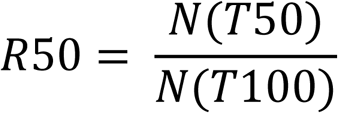

*N(T)* defines the number of unique *aa* sequences that accounts for *T* template frequency.

R50 was calculated from CD3⁺ donor repertoires or, when unavailable, from combined CD4⁺/CD8⁺ pseudo-CD3 repertoires. CD3⁺ donor samples without matched CFSElo (C150, C173) were retained for donor-level analyses but excluded from CFSElo-based comparisons.

Overall TCR CDR3β repertoire diversity was also quantified using the Shannon index and Simpson’s index, while Simpson’s clonality was calculated using published definitions^57^.

### Shared TCRβ Clonotypes

Publicly shared TCRβ clonotypes were derived from the discovery cohort of 20 donors and CFSElo from MLR. A clonotype is shared if it appears in at least 2 donor data points. In total, 114,026 unique clonotype sequences were found to be shared.

### Cross-reactive TCRβ Clonotypes

Cross-reactive TCRβ data set contained *aa* sequences obtained from the public VDJdb database (https://vdjdb.cdr3.net/search, accessed on 01/14/2026), where human EBV, CMV, and Influenza A/B CDR3β *aa* sequences which recognized at least 2 unique epitopes were considered cross-reactive^25^. In addition, cross-reactive sequences were also obtained from a published source examining cross-reactivity within shared CDR3β sequences^24^. In total, our cross-reactive TCRβ set contained 3,797 unique sequences.

### Anti-pathogen TCRβ Clonotypes

26,843 Epstein-Barr Virus (EBV) clonotypes, 5,888 Cytomegalovirus (CMV) clonotypes, and 6,830 Influenza A/B clonotypes were obtained from the public VDJdb data base (https://vdjdb.cdr3.net/search, accessed on 01/14/2026)^25^.

### Statistics

Significance was defined as a *p*-value of 0.05 in all figures unless otherwise specified. Welch’s *t*-tests were used for comparisons of CDR3β length distribution and biochemical features between alloreactive and nonalloreactive clonotypes. Mann-Whitney U tests were used for comparisons of alloreactive unique clonotype fraction and R50 distributions. Fisher’s exact test with odds ratio was used to compare alloreactive clonotype overlap within shared and cross-reactive clonotype sets. 2-way ANOVA was used for decile stratification analyses of donor shared clonotype fraction, mean CDR3β length, and alloreactive clonotype fraction.

### Template Count Stratification

Alloreactive clonotypes in the CFSElo fraction were ranked by template count, defined as the CFSElo frequency normalized to the minimum observed CFSElo frequency within each dataset. Clonotypes were stratified into tertiles, and >2 versus ≤2 template groups, and mean CDR3β length was compared across strata.

### R50 Derivation and Clinical Validation

The optimal RXX threshold was derived using the Area Under the Curve from Receiver Operating Characteristic analysis, with the optimal sensitivity/specificity cutoff determined by the Youden Index. Cumulative Incidence curves were used to analyze time to GVHD in donor populations separated by the R50 cutoff. To evaluate the independent predictive value of R50, multivariable Cox regression models were applied for three endpoints: clinically significant acute GVHD (grade 2-4) within 3 months of transplant, disease relapse within one year, and mortality within one year. Each model was adjusted for donor age, recipient age, HLA matching, donor-recipient sex match status, conditioning regimen, PTCy use, patient disease type, and donor CMV status, with a two-sided significance threshold of 0.05.

### VJ Gene Pairing Analysis

Fisher’s exact test was applied to all unique viable VJ pairing fold-changes between comparison populations, with *p*-values adjusted using the Benjamini-Hochberg method at a False Discovery Rate of *q* = 0.005 to ensure high-confidence detection of significant VJ pairing.

### Biochemical Property Analysis

Biochemical properties of amino acid residues were calculated for hydrophobicity (Kyte-Doolittle scale), charge (formal side-chain charge at physiological pH: +1 for R/K, −1 for D/E, 0 otherwise), polarity (binary polar/nonpolar classification), volume (van der Waals side-chain volume in ų), surface propensity (Janin accessible surface area propensity), flexibility (Bhaskaran-Ponnuswamy normalized B-factor index), and conformational entropy (disorder-promoting amino acid score) as commonly used or previously described for immune receptor sequences^27,30,31^. Each entry point was the mean of a clonotype’s position 6 and position 7 amino acid values, and mean biochemical values for founder alloreactive and all alloreactive clonotypes were compared with nonalloreactive clonotypes using Welch’s *t*-test with Benjamini-Hochberg FDR correction (*q* = 0.05). To statistically validate positional amino acid composition differences, per-position chi-squared tests were performed on 20×2 contingency tables (20 amino acids × 2 groups) across all CDR3β positions, with Benjamini-Hochberg FDR correction (*q* = 0.05) applied across all positions within each comparison.

### Software

Data stratification, processing, and statistical analysis were performed in R-Studio; graphical representation and statistical analysis was performed in GraphPad Prism. Graphical representation was also performed in BioRender.

## Data and Code Availability

All data generated in this study are included in this published article and its supplementary information. Raw TCR sequencing data will be deposited in the NCBI Gene Expression Omnibus (GEO) upon publication. Raw and processed clonotype input files used for downstream analysis are accessible at: https://drive.google.com/drive/u/0/folders/1jRSX3-W59lu3AaqE_XUIciDVZmtWmyXu.

Code for data processing and statistical analysis is available as Supplementary Software at https://github.com/azizilab/GVHD_MLR_Reproducibility/tree/main/MLR_analysis.

Any additional information required to reanalyze the data reported in this paper is available from the lead contact upon request.

## Acknowledgements

We thank the following cores and resources for their support: the Columbia University Center for Translational Immunology Biobank Core and the Flow Cytometry Core. We gratefully acknowledge the Institutional Review Boards of Columbia University, the University of Pennsylvania, and the National Marrow Donor Program as well as all patients and donors who generously consented to participate in this study. We are grateful for support from the Research Stabilization Grant at Columbia University. We are also thankful to Susan DeWolf for her helpful comments and discussions.

## Author Information

X.K.W and R.R., conceived the study. X.K.W., A.U., and K.B. developed MLR protocols and X.K.W., A.U., D.W.H, and K.B. performed sample processing and data collection. D.L.P., E.O.H., A.W.L., and C.A.G provided samples and patient clinical data. X.K.W., A.U., R.M., D.W.H., L.S., M.P., S.C., E.A., and R.R. analyzed the data. X.K.W., A.U., R.M., D.W.H., L.S., M.P., S.C., C.A.G., J.F., E.O.H, A.W.L., D.L.P., M.Y.M., M.S., E.A., and R.R. interpreted the data. X.K.W, A.U., L.S., E.A., and R.R. wrote the manuscript.

