## Supplementary Material for "A Donor T-Cell Receptor Structural Signature Determines Alloreactive Potential and Predicts Acute Graft-Versus-Host Disease"

### Supplemental Material

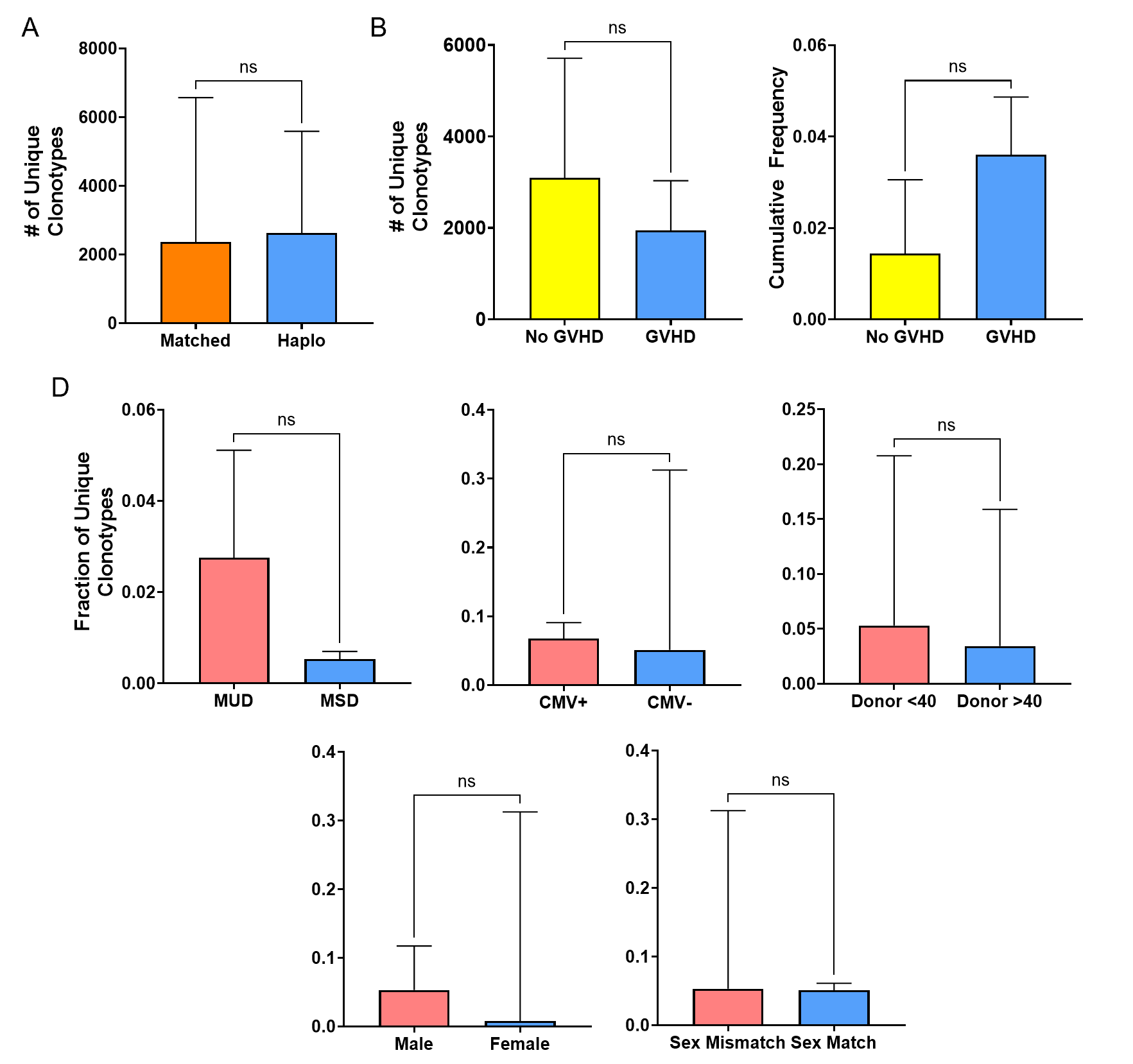

**Supplemental Fig. 1: A)** The median number of alloreactive clonotypes in HLA-mismatched haploidentical donors vs matched donors (combined matched related and unrelated), p = ns. **B)** The median number of unique alloreactive clonotypes in the donor T-cell repertoire comparing patients with and without acute GVHD (grade 2-4) p = ns. **C)** Cumulative alloreactive frequency (Mean ±SEM) of pre-transplant donor repertoire compared in donors of patients with and without acute GVHD (grade 2-4). Welch’s *t*-test used for comparison. **D)** There is no quantitative difference in median fraction of unique alloreactive clonotypes in matched unrelated donors (n = 5) vs matched sibling donors (n = 2) (Median 0.0275 vs 0.0054), CMV+ donors (n = 9) vs CMV- donors (n = 11) (Median 0.0568 vs 0.0511), Donor Age <40 (n = 13) vs Donor Age >40 (n = 7) (Median 0.0530 vs 0.0344), Male donors (n = 16) vs female donors (n = 4) (Median 0.0531 vs 0.0079), Donors sex mismatched with their recipients (n = 11) vs donors without recipient sex mismatch (n = 9) ( Median 0.0530 vs 0.0511). Mann-Whitney U test used for comparison, error bars depict the 95% confidence intervals.

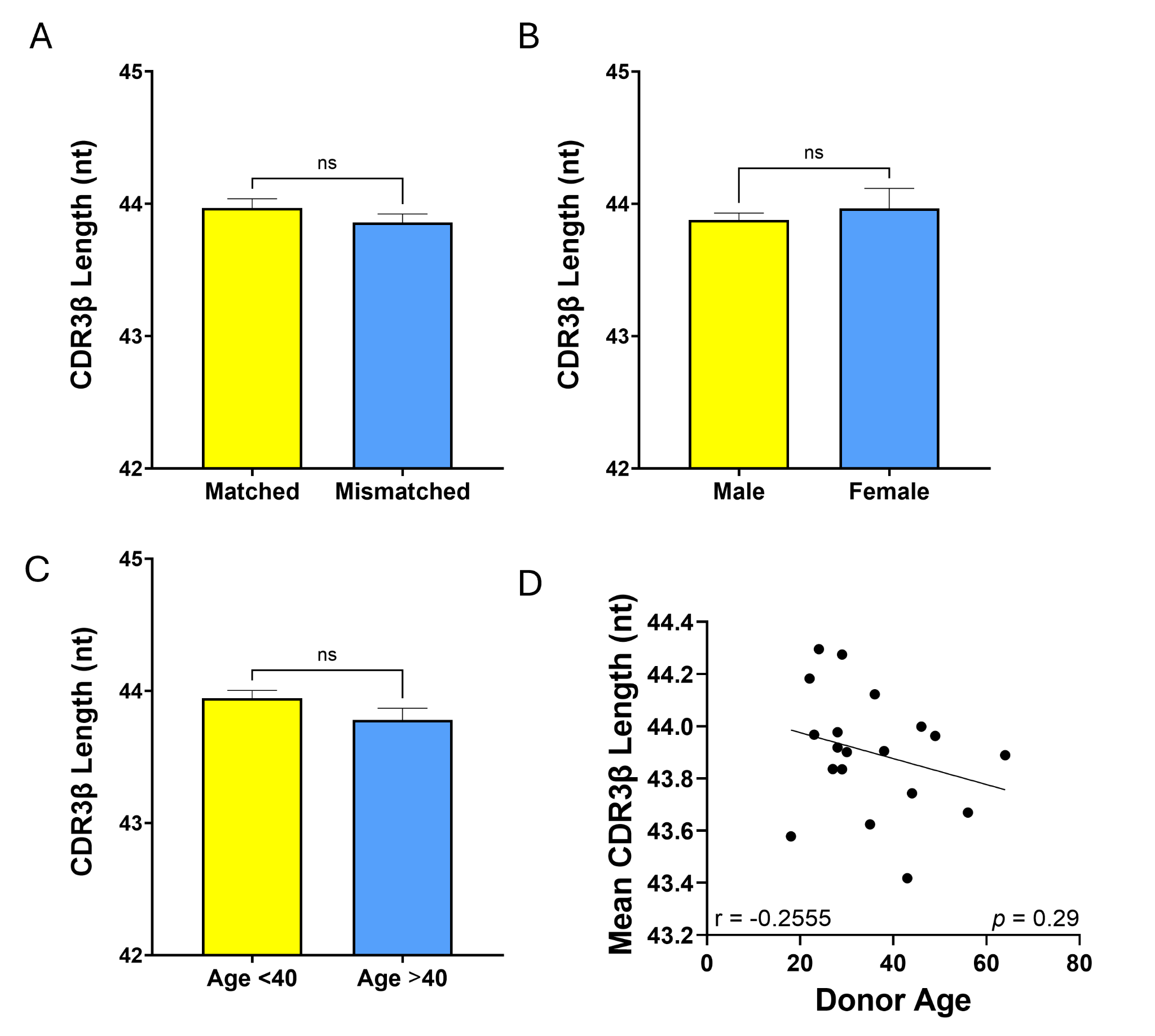

**Supplemental Fig. 2: A-C)** No significant differences in mean CDR3β length of unique donor repertoire clonotypes based on HLA matching, donor sex or age groups. Welch’s *t-*test used to determine significance. **D)** There is no correlation between donor age and mean CDR3β length. Spearman correlation used: r = 0.2514, *p* = 0.29, n = 20.

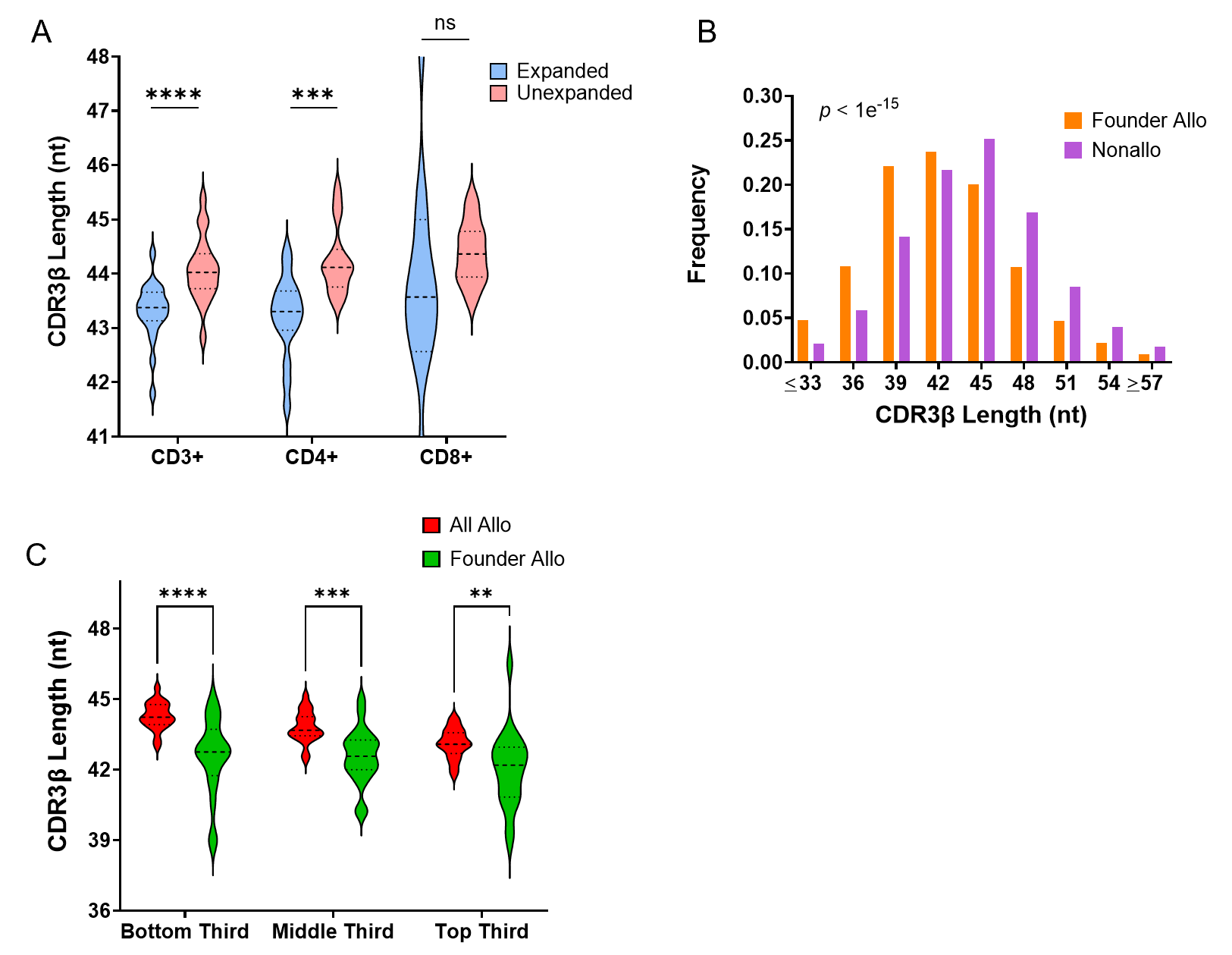

**Supplemental Fig 3:** **A)** Violin plots of CDR3β length distributions of alloreactive clonotypes stratified into expanded (>2 templates) and unexpanded (≤ 2 templates) subpopulations within the CFSElo fraction. Statistical significance was determined using a Mann-Whitney U test. **B)** CDR3β length distribution of founder alloreactive vs nonalloreactive clonotypes. Welch’s *t-*test used to determine significance. **C)** Violin plots comparing CDR3β length distributions of founder alloreactive clones (measurable in a donor baseline blood sample) vs. all alloreactive clonotypes, stratified by their frequencies in the CFSElo fraction. Mann-Whitney U test used to determine significance ******p* < 0.05, ***p* < 0.01, ****p* < 0.001, *****p* < 0.0001.

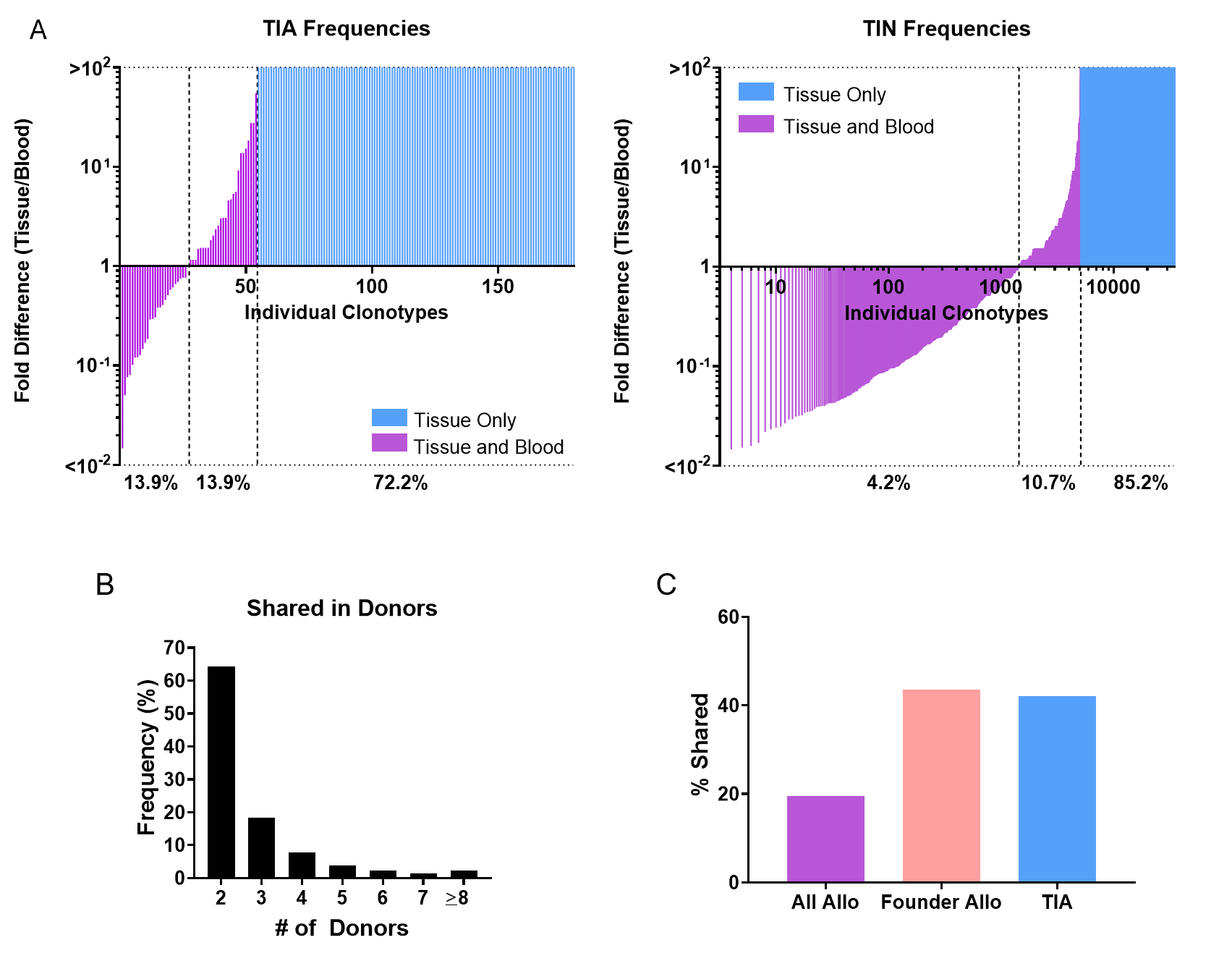

**Supplemental Fig. 4:** **A)** Distribution of repertoire frequency ratio between recipient tissue and PBMC for tissue-infiltrating alloreactive (TIA) clonotypes. 72.2% of TIA do not have a detectable PBMC counterpart and thus the tissue/PBMC ratio is infinite. Distribution of tissue-infiltrating nonalloreactive (TIN) clonotypes. **B)** Bar plots comparing the frequencies by which shared clonotypes appear in multiple donor repertoires (grouped by number of donors). n = 114,026. **C)** Percentage of unique clonotypes that are shared in various alloreactive categories.

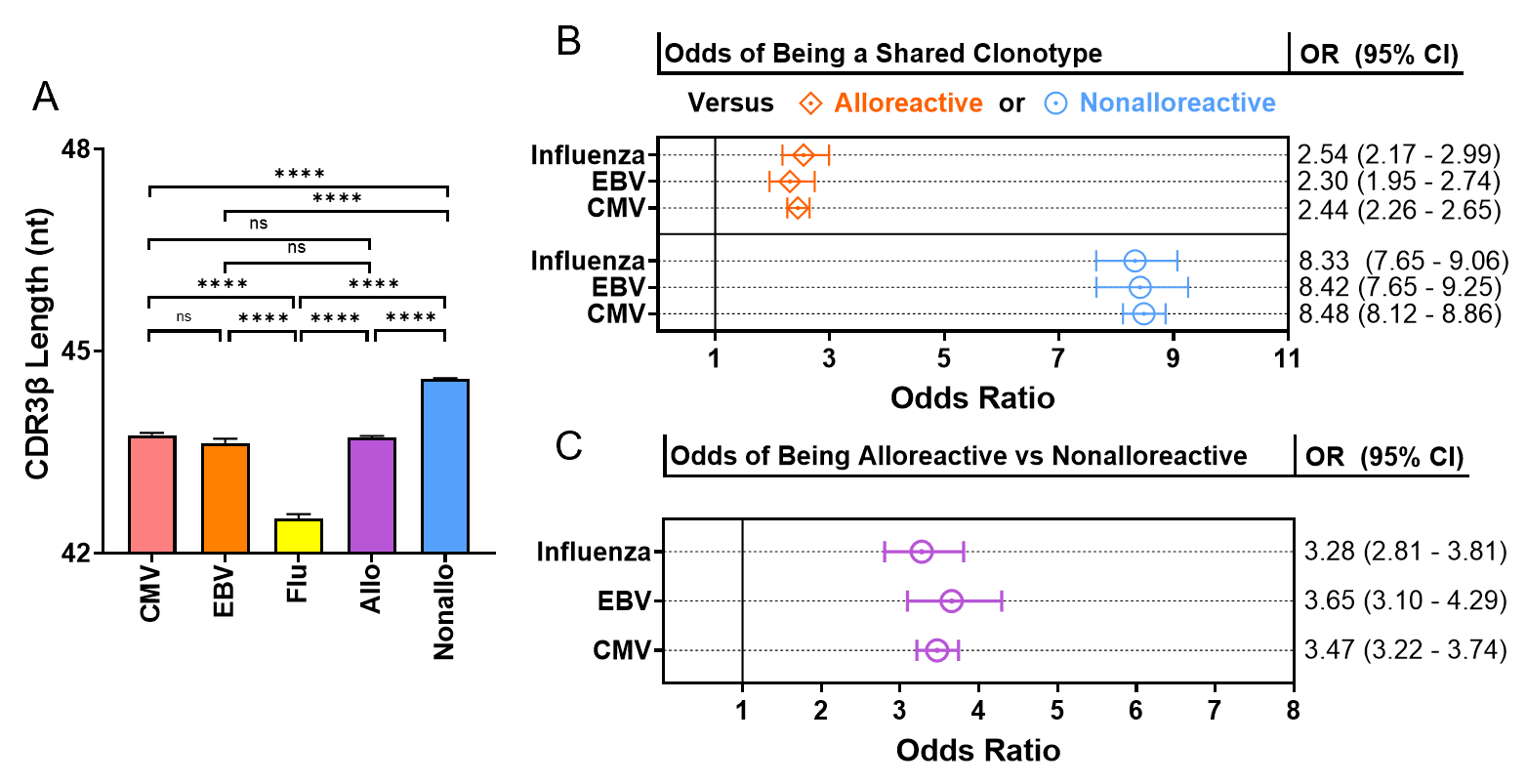

**Supplemental Fig. 5: A)** CDR3ꞵ lengths (mean ± SEM) of publicly available viral antigen-recognizing clonotypes for CMV, EBV, Influenza (A and B) along with comparison to all MLR-defined alloreactive and nonalloreactive clonotypes. ******p* < 0.05, ***p* < 0.01, ****p* < 0.001, *****p* < 0.0001. Welch’s *t-*test. **B)** Odds ratio (95% CI) of publicly available viral anti-pathogen clonotypes being shared in comparison to MLR-defined nonalloreactive clonotypes. Odds ratio with 95% CI of publicly available viral anti-pathogen clonotypes being shared in comparison to alloreactive or nonalloreactive clonotypes. **C)** Odds ratio (95% CI) of publicly available viral anti-pathogen clonotypes of being alloreactive in comparison to nonalloreactive clonotypes.

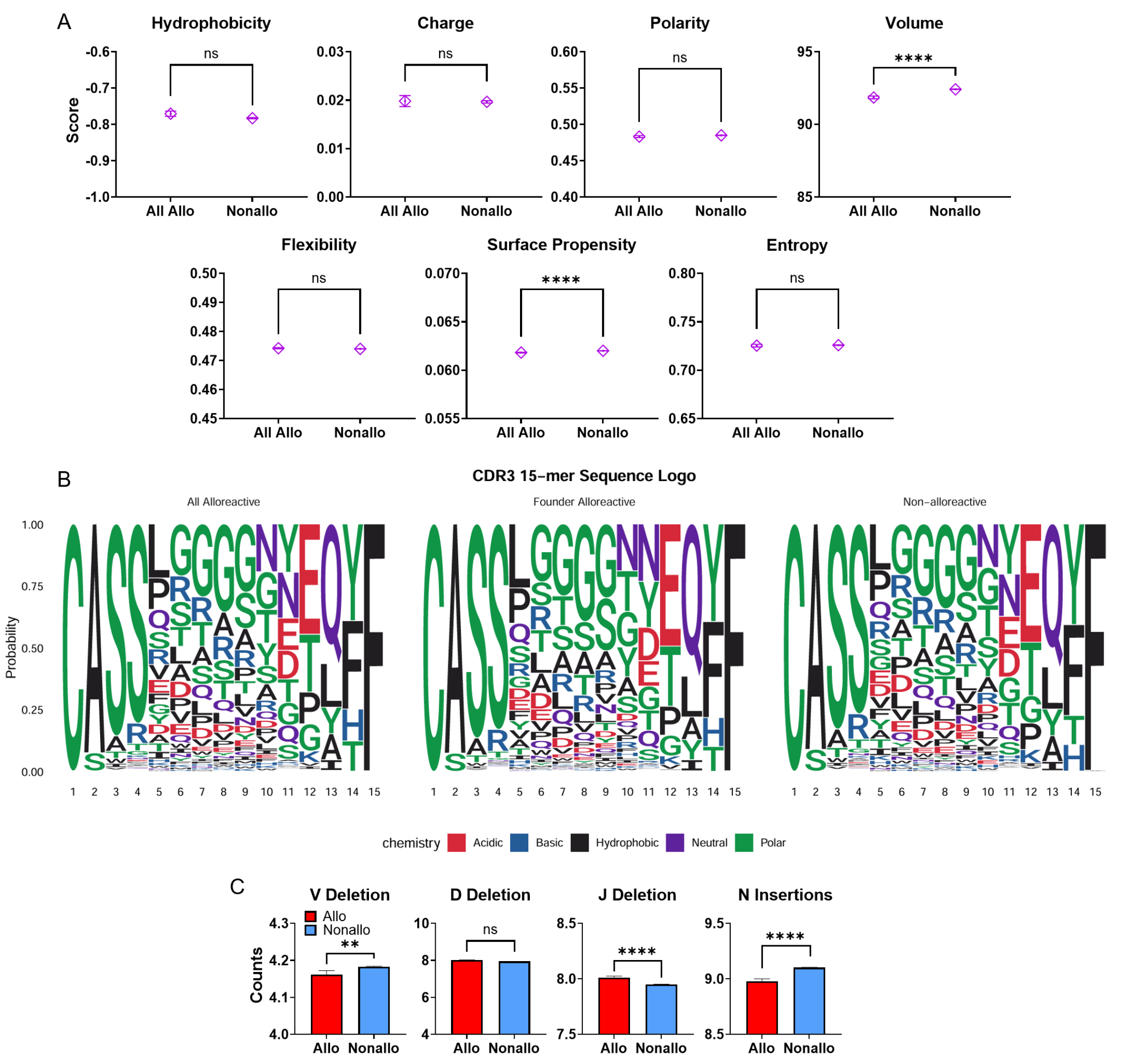

**Supplemental Fig. 6: A)** Calculated mean (±SEM) of hydrophobicity (Kyte-Doolittle scale), charge, polarity, volume, flexibility, surface propensity, and entropy for amino acid residues in position 6 and position 7 of CDR3β chain for all alloreactive vs nonalloreactive clonotype groups **B)** Logos depict amino acid frequencies across 15-mer CDR3ꞵ regions for all alloreactive, founder alloreactive, and nonalloreactive groups. Colors indicate amino acid chemistry. Per-position chi-squared analysis confirmed significantly distinct amino acid composition at positions 6 and 7 between founder alloreactive and nonalloreactive clonotypes. FDR (*q* = 0.05) adjusted *p* = 1.46e^-4^ for both positions. **C)** Mean (±SEM) counts of V deletions, D deletions, J deletions, and N insertions in the CDR3β region in all alloreactive vs nonalloreactive clonotypes. **p* < 0.05, ***p* < 0.01, ****p* < 0.001, *****p* < 0.0001. Welch’s *t*-test used for all comparisons.

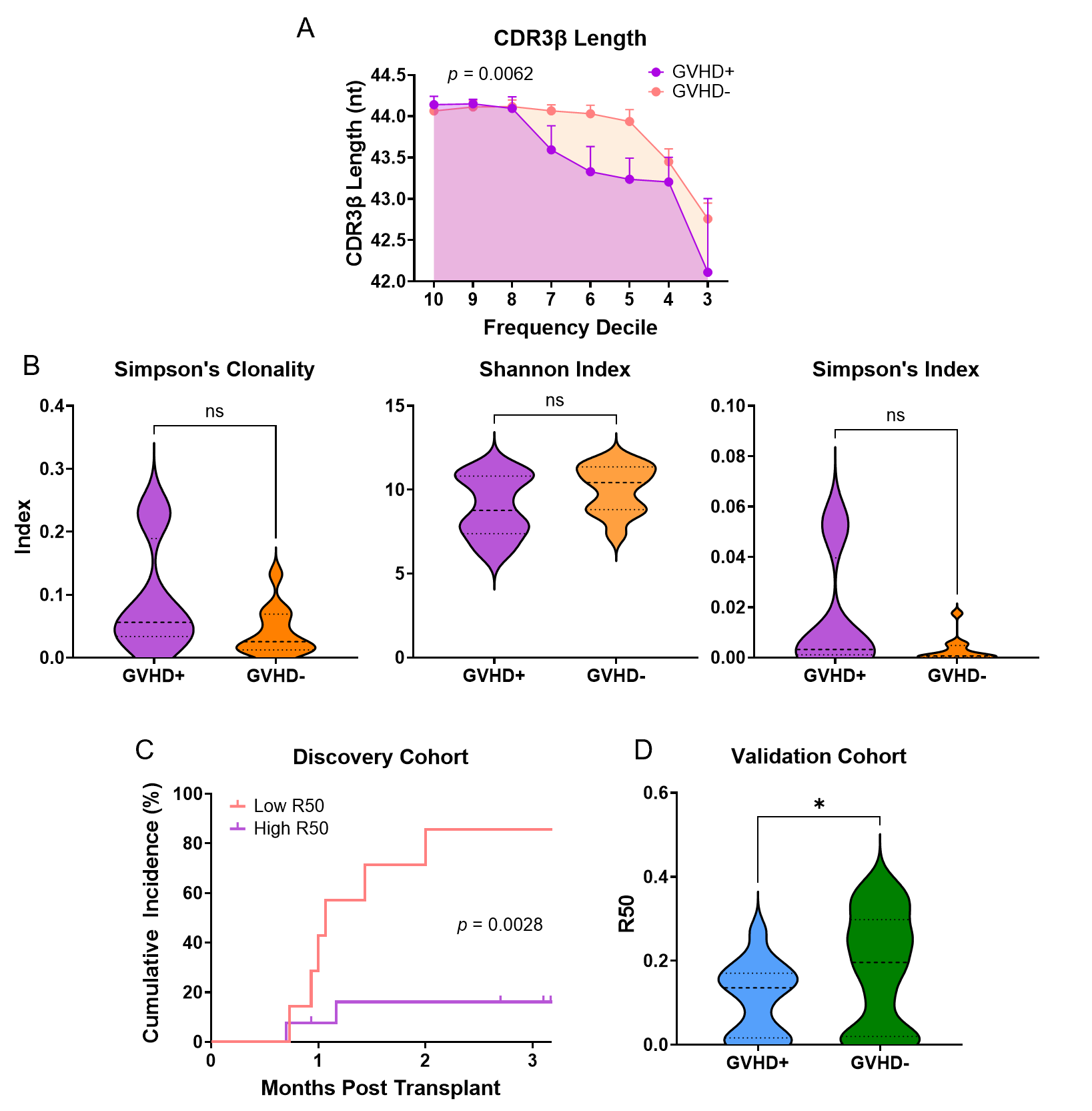

**Supplemental Fig. 7: A)** Comparison of mean (±SEM) CDR3ꞵ length in deciles for patients with and without acute GVHD in the discovery cohort. 2-way ANOVA test used to determine statistical significance. **B)** Conventional donor repertoire diversity metrics (Simpson's clonality, Shannon entropy, and Simpson's index) did not significantly differ between GVHD+ and GVHD- donor groups. Mann-Whitney U test used. **C)** Cumulative incidence plots of time to acute GVHD for the discovery cohort (n = 20), stratified by the derived optimal diagnostic cutoff R50 (< 0.1714). **D)** Median donor R50 with 95% CI for GVHD+ and GVHD- patients in the validation cohort (n = 57). *p* = 0.043. Mann-Whitney U test used.

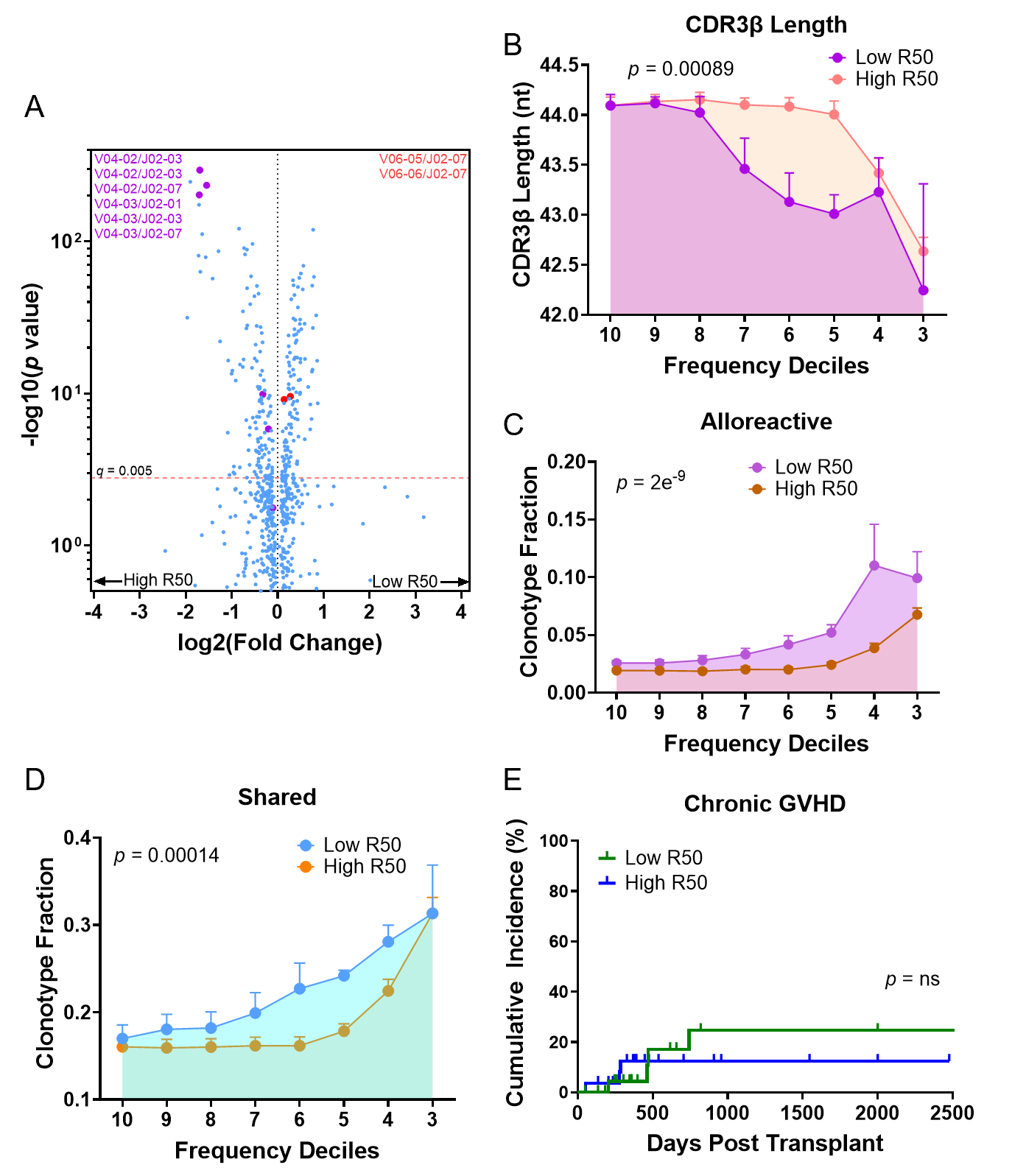

**Supplemental Fig. 8: A**) Volcano plot showing VJ gene combination usage in Low R50 vs. High R50 clonotypes from the discovery cohort . Fisher’s exact test used to calculate the significance of the differences between the groups. Benjamini-Hochberg test using a *q* value of 0.005 was used to calculate the cut-off for significance. Low R50 VJ pairings that significantly correlated with GVHD+/Allo-biased status are listed. Red values denote VJ pairings that are GVHD+/Allo-biased and purple values denote GVHD-/Nonallo-biased VJ pairings. **B)** Comparison of average CDR3β clonotype length in diversity deciles intervals for patients who have Low R50 scoring donors (n = 7) and those who have High R50 scoring donors (n = 13). **C)** Comparison of Low R50 donors vs High R50 donors with greater alloreactivity and **D)** fraction of shared clonotypes by decile frequency intervals. 2-way ANOVA test used for all with *p*-value of R50 grouping. E) Cumulative incidence plots of time to moderate-severe chronic GVHD for the validation cohort (n = 57), separated using the derived optimal diagnostic cutoff R50 (< 0.1714).

| **Supplemental Table 1.** | |
| --- | --- |
| **Number of HCT Patients, *n*** | 57 |
| **Institution** |  |
| Columbia University Medical Center, *n* | 23 |
| University of Pennsylvania, *n* | 34 |
| **Donor Sex** |  |
| Male, *n* | 36 |
| Female, *n* | 19 |
| Unknown, *n* | 2 |
| **Recipient Sex** |  |
| Male, *n* | 28 |
| Female, *n* | 27 |
| Unknown, *n* | 2 |
| **Donor/Recipient Sex** |  |
| Matched, *n* | 33 |
| Mismatched, *n* | 22 |
| Unknown, *n* | 2 |
| **Donor Age, Median (Range)** | **38 (18-70)** |
| ≥40, *n* | 30 |
| <40, *n* | 27 |
| **Transplant Recipient Age, Median (Range)** | **59 (21-72)** |
| ≥60, *n* | 25 |
| <60, *n* | 28 |
| Unknown, *n* | 4 |
| **Donor CMV Status** |  |
| Yes, *n* | 23 |
| No, *n* | 32 |
| Unknown, *n* | 2 |
| **Acute GVHD Grade (< 3 Months Post-Transplant)** |  |
| Not Significant (Grade 0-1), *n* | 43 |
| Significant (Grade 2-4), *n* | 14 |
| **Chronic GVHD State** |  |
| None-Mild, *n* | 46 |
| Moderate-Severe, *n* | 7 |
| Unknown, *n* | 4 |
| **Disease Diagnosis** |  |
| Acute Myeloid Leukemia, *n* | 32 |
| Acute Lymphoid Leukemia, *n* | 1 |
| Aplastic Anemia, *n* | 1 |
| Chronic Myeloid Leukemia, *n* | 1 |
| Chronic Myelomonocytic Leukemia, *n* | 1 |
| Myelofibrosis, *n* | 2 |
| Myelodysplastic Syndrome, *n* | 6 |
| Hodgkin Lymphoma, *n* | 3 |
| Non-Hodgkin Lymphoma, *n* | 4 |
| Cutaneous T-cell Lymphoma, *n* | 3 |
| T-Cell Acute Lymphoblastic Leukemia, *n* | 1 |
| T-Cell Prolymphocytic Leukemia, *n* | 1 |
| Sickle Cell Disease, *n* | 1 |
| **Donor Type** |  |
| Matched Unrelated Donor (10/10 HLA), *n* | 28 |
| Mismatched Unrelated Donor (9/10 HLA), *n* | 1 |
| Matched Sibling Donor (10/10 HLA), *n* | 11 |
| Other Matched Related Donor (10/10 HLA), *n* | 3 |
| Haploidentical, *n* | 12 |
| Unknown, *n* | 2 |
| **Post-Transplant Cyclophosphamide** |  |
| Yes, *n* | 14 |
| No, *n* | 43 |
| Unknown, *n* | 1 |
| **Conditioning Regimen** |  |
| Myeloablative, *n* | 12 |
| Reduced Intensity, *n* | 45 |

**Supplemental Table 1. Patient and Donor Characteristics in the Expanded (Validation) Cohort (n = 57)**

| **Supplemental Table 2.** | | | | |
| --- | --- | --- | --- | --- |
| **Patient ID** | **Tissue Sources** | **Biopsy - Days PT** | **aGVHD Grade 2-4 Status** | **TIA Clonotypes, *n*** |
| C051 | Colon | 39 | No | 0 |
| C067 | Colon, Duodenum, Gastric Body | 23 | Yes | 11 |
| C069 | Duodenum, Esophagus, Stomach | 23 | Yes | 8 |
| C070 | Combined Upper and Lower Gut | 93 | No | 5 |
| C081 | Upper GI, Lower GI | 208 | No | 105 |
| C086 | Upper GI, Lower GI, Skin | 140 and 240 | Yes | 49 |
| C101 | Lower GI | 30 | Yes | 5 |

**Supplemental Table 2. Post-Transplant Tissues Collected at time of suspected GVHD**. PT = Post-transplant, TIA = Tissue-infiltrating alloreactive, aGVHD = acute GVHD.

| **Supplemental Table 3.** | | | | | |
| --- | --- | --- | --- | --- | --- |
| **RXX** | ***R10*** | ***R30*** | ***R50*** | ***R70*** | ***R90*** |
| **AUC** | 0.7292 | 0.8542 | 0.8958 | 0.875 | 0.875 |
| **Youden Index Cutoff** | 0.0002755 | 0.04983 | 0.1714 | 0.566 | 0.8553 |
| **Sensitivity** | 0.75 | 1 | 0.75 | 1 | 1 |
| **Specificity** | 0.6667 | 0.6667 | 0.9167 | 0.6667 | 0.6667 |
| **Likelihood Ratio** | 2.25 | 3 | 9 | 3 | 3 |
| **PPV** | 0.4909341 | 0.5625246 | 0.7941831 | 0.5625246 | 0.5625246 |
| **NPV** | 0.8615444 | 1 | 0.8953522 | 1 | 1 |

**Supplemental Table 3:** **RXX and statistical donor clonotype diversity cutoffs for GVHD prediction.** PPV and NPV are calculated from grade 2-4 acute GVHD prevalence of 24%, derived from the validation cohort. AUC = Area under the curve for Receiver Operator Characteristic (ROC) curve. PPV = Positive Predictive Value and NPV = Negative Predictive Value.

| **Supplemental Table 4. Acute GVHD - Univariate** | | | |
| --- | --- | --- | --- |
| **Variable** | **Hazard Ratio** | **Confidence Interval (95%)** | ***p* - value** |
| *R50 (< 0.1714 vs ≥ 0.1714)* | **4.059** | 1.266 to 17.96 | **0.0170*** |
| *Donor Age (≥40 vs <40)* | 0.8105 | 0.2667 to 2.332 | 0.6962 |
| *Donor Recipient Sex Mismatch/Match* | 2.342 | 0.8138 to 7.121 | 0.1135 |
| *Donor CMV Status (+/-)* | 0.7872 | 0.2418 to 2.280 | 0.6648 |
| *Myeloid/Non-Myeloid Disease* | 1.162 | 0.3627 to 5.139 | 0.8151 |
| *Recipient Age (>60 vs ≤ 60)* | 0.5129 | 0.1368 to 1.630 | 0.2622 |
| *PTCy GVHD Prophylaxis (+/-)* | 2.743 | 0.9005 to 7.913 | 0.0740 |
| *Myeloablative/Reduced Intensity Conditioning Regimen* | 2.171 | 0.6664 to 6.290 | 0.1851 |
| *HLA Mismatched/Matched* | 3.040 | 0.9972 to 8.779 | 0.0505 |

**Supplemental Table 4: Univariate Analysis for R50 as a predictor of acute GVHD**. Data of 57 transplant donor-recipients in the expanded cohort. GVHD was defined as grade 2-4 acute GVHD within 3 months of HCT. PTCy = Post-transplant Cyclophosphamide. Cox regression analysis used to calculate hazard ratios for acute GVHD within 3 months post-transplant. **p* < 0.05

| **Supplemental Table 5. Acute GVHD - Multivariate** | | | |
| --- | --- | --- | --- |
| **Variable** | **Hazard Ratio** | **Confidence Interval (95%)** | ***p* - value** |
| *R50 (< 0.1714 vs ≥ 0.1714)* | **6.366** | 1.573 to 33.60 | **0.0087**** |
| *Donor Age (≥40* *vs <40)* | 0.6321 | 0.1588 to 2.515 | 0.5066 |
| *Donor Recipient Sex Mismatch/Match* | 1.581 | 0.3988 to 5.894 | 0.4981 |
| *Donor CMV Status (+/-)* | 1.527 | 0.3581 to 6.483 | 0.5594 |
| *Myeloid Diseases/Non-Myeloid* | 1.706 | 0.2255 to 18.20 | 0.6188 |
| *Recipient Age (>60 vs ≤ 60)* | 0.8598 | 0.1235 to 5.737 | 0.8754 |
| *PTCy GVHD Prophylaxis (+/-)* | 1.025 | 0.02710 to 20.09 | 0.9883 |
| *Myeloablative/Reduced Intensity Conditioning Regimen* | 1.06 | 0.1583 to 7.116 | 0.9515 |
| *HLA Mismatch/Matched* | 4.86 | 0.2270 to 127.7 | 0.3286 |

**Supplemental Table 5. Multivariate Analysis for R50 as a predictor of acute GVHD***.* Data for 57 transplant donor-recipients in the expanded cohort. GVHD was defined as grade 2-4 acute GVHD within 3 months of HCT. PTCy = Post-transplant Cyclophosphamide. Multivariate Cox regression analysis used to determine hazard ratios for acute GVHD. ***p* < 0.01

| **Supplemental Table 6. Disease Relapse** | | | |
| --- | --- | --- | --- |
| **Variable** | **Hazard Ratio** | **Confidence Interval (95%)** | ***p* - value** |
| *R50 (< 0.1714 vs ≥ 0.1714)* | 1.428 | 0.3354 to 6.921 | 0.6361 |
| *GVHD Status (+/-)* | 1.154 | 0.1310 to 6.074 | 0.8810 |
| *Donor Age (≥40 vs <40)* | 0.7347 | 0.1513 to 3.241 | 0.6875 |
| *Donor Recipient Sex Mismatch/Match* | 0.4657 | 0.09169 to 1.774 | 0.2776 |
| *Donor CMV Status (+/-)* | 0.4763 | 0.09679 to 1.900 | 0.3034 |
| *Myeloid/Non-Myeloid Disease* | **0.1679** | 0.03590 to 0.7011 | **0.0145*** |
| *Recipient Age (>60 vs ≤ 60)* | 0.9295 | 0.2217 to 3.761 | 0.9177 |
| *PTCy GVHD Prophylaxis (+/-)* | 1.286 | 0.1987 to 7.205 | 0.7782 |
| *Myeloablative/Reduced Intensity Conditioning Regimen* | 0.3957 | 0.01975 to 2.742 | 0.3776 |
| *HLA Mismatch/Match* | 0.4672 | 0.04656 to 3.277 | 0.4554 |

**Supplemental Table 6: Multivariate Analysis for R50 as a predictor of Disease Relapse within a Year**. Data for 57 transplant donor-recipients in the expanded cohort. GVHD was defined as Grade 2-4 acute GVHD within 3 months of HCT. Relapse of disease defined as occurring within a year of HCT. PTCy = Post-transplant Cyclophosphamide. Multivariate Cox regression analysis used to calculate hazard ratios for disease relapse. **p* < 0.05

| **Supplemental Table 7. 1-Year Mortality** | | | |
| --- | --- | --- | --- |
| **Variable** | **Hazard Ratio** | **Confidence Interval (95%)** | ***p* - value** |
| *R50 (< 0.1714 vs ≥ 0.1714)* | 0.6911 | 0.1400 to 3.574 | 0.6465 |
| *GVHD Status (+/-)* | **12.81** | 2.272 to 81.78 | **0.0045**** |
| *Relapse Status (+/-)* | **6.784** | 1.745 to 31.57 | **0.0056**** |
| *Donor Age (≥40 vs <40)* | 3.388 | 0.6045 to 20.71 | 0.1640 |
| *Donor Recipient Sex Mismatch/Match* | 0.6929 | 0.1112 to 3.255 | 0.6507 |
| *Donor CMV Status (+/-)* | 0.6117 | 0.1098 to 3.054 | 0.5513 |
| *Myeloid/Non-Myeloid Disease* | 0.3741 | 0.05434 to 2.448 | 0.3025 |
| *Recipient Age (>60 vs ≤ 60)* | 2.281 | 0.3418 to 19.46 | 0.4035 |
| *PTCy GVHD Prophylaxis (+/-)* | 3.427 | 0.1488 to 42.59 | 0.3935 |
| *Myeloablative/Reduced Intensity Conditioning Regimen* | 2.327 | 0.2551 to 28.84 | 0.4574 |
| *HLA Mismatch/Match* | 0.08468 | 0.002460 to 3.113 | 0.1728 |

**Supplemental Table 7: Multivariate Analysis for R50 as a predictor of 1-Year Mortality**. Data for 57 transplant donor-recipients in the expanded cohort. GVHD was defined as grade 2-4 acute GVHD within 3 months of HCT. Relapse of disease defined as occurring within a year of HCT. PTCy = Post-transplant Cyclophosphamide. Multivariate Cox regression analysis conducted for 1-year mortality. **p* < 0.05, ***p* < 0.01
